# Multiplex generation and single cell analysis of structural variants in a mammalian genome

**DOI:** 10.1101/2024.01.22.576756

**Authors:** Sudarshan Pinglay, Jean-Benoit Lalanne, Riza M. Daza, Jonas Koeppel, Xiaoyi Li, David S. Lee, Jay Shendure

## Abstract

The functional consequences of structural variants (SVs) in mammalian genomes are challenging to study. This is due to several factors, including: 1) their numerical paucity relative to other forms of standing genetic variation such as single nucleotide variants (SNVs) and short insertions or deletions (indels); 2) the fact that a single SV can involve and potentially impact the function of more than one gene and/or *cis* regulatory element; and 3) the relative immaturity of methods to generate and map SVs, either randomly or in targeted fashion, in *in vitro* or *in vivo* model systems. Towards addressing these challenges, we developed *Genome-Shuffle-seq*, a straightforward method that enables the multiplex generation and mapping of several major forms of SVs (deletions, inversions, translocations) throughout a mammalian genome. *Genome-Shuffle-seq* is based on the integration of “shuffle cassettes’’ to the genome, wherein each shuffle cassette contains components that facilitate its site-specific recombination (SSR) with other integrated shuffle cassettes (via Cre-loxP), its mapping to a specific genomic location (via T7-mediated *in vitro* transcription or IVT), and its identification in single-cell RNA-seq (scRNA-seq) data (via T7-mediated *in situ* transcription or IST). In this proof-of-concept, we apply *Genome-Shuffle-seq* to induce and map thousands of genomic SVs in mouse embryonic stem cells (mESCs) in a single experiment. Induced SVs are rapidly depleted from the cellular population over time, possibly due to Cre-mediated toxicity and/or negative selection on the rearrangements themselves. Leveraging T7 IST of barcodes whose positions are already mapped, we further demonstrate that we can efficiently genotype which SVs are present in association with each of many single cell transcriptomes in scRNA-seq data. Finally, preliminary evidence suggests our method may be a powerful means of generating extrachromosomal circular DNAs (ecDNAs). Looking forward, we anticipate that *Genome-Shuffle-seq* may be broadly useful for the systematic exploration of the functional consequences of SVs on gene expression, the chromatin landscape, and 3D nuclear architecture. We further anticipate potential uses for *in vitro* modeling of ecDNAs, as well as in paving the path to a minimal mammalian genome.

## Introduction

Major classes of human genetic variation include SNVs, indels, simple sequence repeat variants, and genomic SVs (*e.g.* deletions, insertions, inversions, and duplications over 50 bp, and chromosomal translocations) (*1*, *2*). Historically, genomic SVs have been at a distinct disadvantage in terms of our ability to study their functional consequences, as compared to SNVs or indels. This is unfortunately the case for not only paradigms of genetic analysis that rely on living humans, but also genetic analyses conducted entirely in *in vitro* or *in vivo* model systems.

For human genetics, *de novo* SVs are over 100-fold less frequent than *de novo* SNVs per human generation (*3*). Therefore, SVs are less likely to recur, and when they do recur, are less likely to do so in a way that permits the unambiguous implication of a specific functional unit (*e.g.* recurrent disruption of identical sets of genes or regulatory elements). This contrasts with the recurrence of disruptive SNVs or indels within a single gene or regulatory element, which routinely allows for the assignment of causality in Mendelian studies. Furthermore, SVs’ lower rate of *de novo* occurrence, together with a greater likelihood of deleterious fitness effects (because SVs disrupt orders-of-magnitude more base-pairs [bp] than SNVs or indels), contribute to an even greater numerical paucity among standing genetic variants in human populations (*3–7*). Far fewer SVs than SNVs or indels reach the common allele frequencies that would allow for the well-powered detection of phenotypic effects by genome-wide association studies (GWAS). To an extent, these limitations can be addressed with larger cohort sizes, but this has limits. For example, although every possible SNV compatible with life is likely present in a living human (*8*), this is unlikely to be the case for all possible SVs.

For laboratory-based genetics, a plethora of strategies have been developed to introduce SNVs or indels into model systems for functional analysis. These include classic chemical mutagenesis screens as well as their modern equivalent, base editing screens (*9*). They also include using massively parallel DNA synthesis or mutagenic PCR to achieve saturation mutagenesis of a sequence of interest, which can then be studied either inside or outside of its native genomic context (*10*, *11*). Although many specific SNVs or indels generated by such methods have yet to be observed in a living human, their analysis can nonetheless be informative in myriad ways, *e.g.* for implicating specific genes in specific phenotypes (*12*), for systematically characterizing the distribution of effect sizes of regulatory or coding variants (*10*, *11*, *13*), for pre-computing the clinical consequences of potential variants in disease-associated genes (*11*), for optimizing immunotherapies (*14*), etc. However, once again SVs are at a clear disadvantage as compared to SNVs or indels, here due to the relative immaturity of methods to generate and map SVs in model systems.

As a consequence of these disadvantages, there remain numerous unanswered “structure-function” questions about the human genome that relate to its properties at the scale of SVs rather than SNVs or indels. Genes, exons and *cis-*regulatory elements are scattered over vast distances, ordered and oriented in specific ways in specific genomes. However, our understanding of the functional implications of these distances, orders and orientations arguably remains very shallow. For example, roughly one-quarter of the human genome is composed of gene deserts (*15*). Although patterns of conservation suggest at least some sequences within such deserts are functional, the deletion of even megabase-sized gene deserts yields viable mice with no discernable phenotype (*16*, *17*). Other non-genic SVs clearly cause Mendelian disorders, contribute to complex disease risk, or underlie evolutionary adaptations (*18*), but discerning *how* they do so in individual cases often remains unclear. Some mammalian genomes differ by over 1 billion bp in size from the human genome (*19*), and even between similarly sized mammalian genomes such as mouse and human, over 1 billion bp have both been gained and lost over evolutionary time via structural variation (*20*). Beyond the germline, somatic SVs of all kinds, including some cancer-specific forms of structural variation like chromothripsis and ecDNAs (*21*, *22*), are well established to play a critical role in the initiation and progression of nearly all human cancers.

Various strategies have been developed for engineering SVs. For example, site-specific recombinase (SSR) recognition sites can be introduced to specific locations, such that their recombination results in a specific SV, or even ecDNA species, of interest (*23–25*). However, this is labor-intensive, and only results in one or a handful of SVs to study. Alternatively, CRISPR/Cas9 can be used to simultaneously drive double-stranded breaks (DSB) at multiple locations, which can result in the generation of SVs, potentially even genome-wide (*26–28*). But CRISPR/Cas9-based SV induction is challenged by inefficiency, imprecision, DSB toxicity, and absence of a means to efficiently map which cells harbor which (if any) induced SVs. Towards a genome-wide SSR-based approach, inspiring work by Sauvageau and colleagues deployed retroviral introduction of SSR recognition sites to generate a panel of mESC clones bearing nested deletions covering ∼25% of the mouse genome (*29*, *30*). However, the method is still limited by the lack of an efficient means of mapping the initial SSR recognition site positions nor for genotyping post-induction SVs. In yeast, chromosome-specific or genome-wide “scrambles” were achieved by first building synthetic chromosomes containing many SSR recognition sites (*31–33*), but for mammalian genomes, whole chromosome or whole genome synthesis remains impractical. Finally, in all cases (including yeast) where a larger number of SVs are (potentially) induced, the recovery, verification and/or quantitation of SVs relies on inefficient or expensive methods (*e.g.* single cell cloning, whole genome sequencing, karyotyping), which markedly limits what can be achieved, particularly for mammalian models.

Motivated by these technical gaps, we developed *Genome-Shuffle-seq*, a straightforward method for multiplex generation of large-scale SVs throughout a mammalian genome (**Fig. 1**). A key attribute of *Genome-Shuffle-seq* is that it facilitates the facile mapping and genotyping of the breakpoints of induced SVs at base-pair resolution within a population of cells. As a proof-of-concept, we apply *Genome-Shuffle-seq* to induce and map thousands of genomic SVs of several major forms (deletions, inversions, chromosomal translocations, ecDNAs) in mESCs, and to map their coordinates at base-pair resolution without any need for whole genome sequencing. We further demonstrate that we can co-capture the identities of induced SVs as part of a single cell transcriptome, laying the foundation for pooled cellular screens of thousands to millions of mammalian SVs.

**Fig 1.**
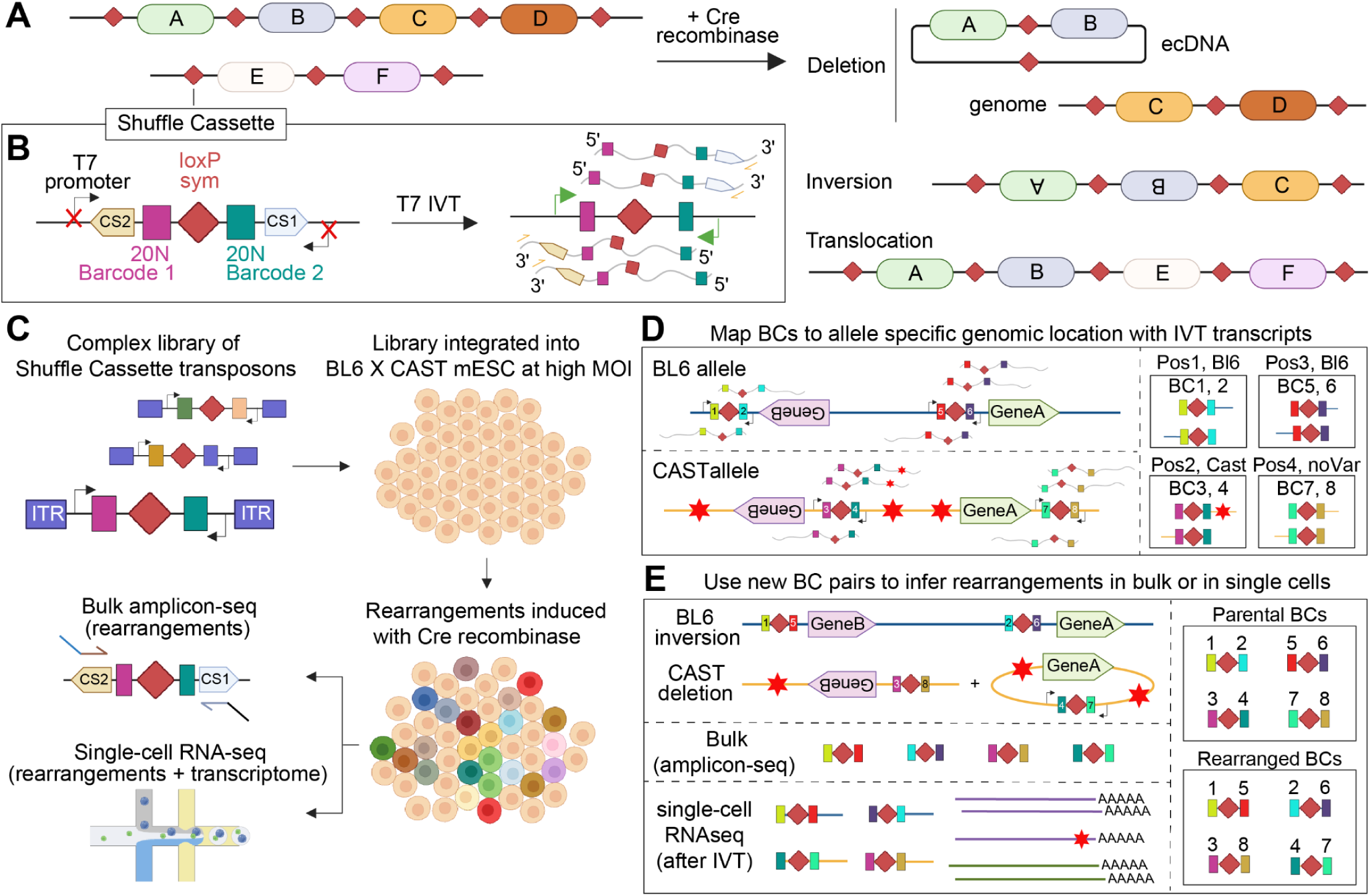
Schematic of *Genome-Shuffle-seq* for the pooled construction and efficient characterization of rearranged mammalian genomes at single-cell resolution. **A)** Arrays of integrated loxPsym sites are recombined by Cre recombinase to yield three classes of SVs. **B)** Schematic of the shuffle cassette, which contains a loxPsym site flanked by unique 20N barcodes, capture sequences for scRNA-seq (CS1, CS2), and phage polymerase promoters that are inert in live mammalian cells but activated upon *in vitro* (IVT) or *in situ* (IST) transcription with T7 polymerase. **C)** Workflow of a *Genome-Shuffle-seq* experiment. **D)** Shuffle cassette insertion sites can be mapped by sequencing T7-derived transcripts from IVT or IST and associating a pair of unique barcodes (numbered 1-8 in schematic) with an allele-specific genomic location. Red stars indicate variants between the BL6 and Castaneus haplotypes in genomic DNA flanking the integration site. **E)** Induced SVs can be inferred by novel barcode combinations that are only observed in amplicons or scRNA-seq data from cells that have been exposed to Cre recombinase. As the genomic coordinates of the parental barcodes are known from IVT-based mapping of their locations in the parental cell population, the identity of the barcodes making up each novel combination is sufficient to infer both the class (deletion, inversion, translocation) as well as the precise genomic coordinates involved in each induced SV.

### Design of Genome-Shuffle-seq

*Genome-Shuffle-seq* is based on the integration of “shuffle cassettes’’ to a mammalian genome (**Fig. 1A-C**). The shuffle cassettes are designed to facilitate the mapping of genomic integration coordinates, the generation of SVs via SSR between pairs of shuffle cassettes, and the efficient recovery of genotype information. The shuffle cassette is 176 bp in length and has four key features (**Fig. 1B**): 1) A loxPsym site capable of being recombined with other loxPsym sites at high efficiency by Cre recombinase. Recombination between two copies of this symmetric variant of the canonical loxP site is expected to lead to both deletions and inversions at roughly equal frequencies (*31*, *34*), as well as translocations; 2) Immediately flanking the loxPsym site, a pair of random 20 nucleotide (nt) barcodes, which uniquely tag each integration of the shuffle cassette, or its recombined derivatives; 3) Further flanking those, a pair of PCR primer binding sites (*35*) (**Fig. S1A**); and 4) At the outer boundaries, a pair of convergently oriented phage T7 RNA polymerase promoters that are inert in mammalian cells but can be activated after fixation for *in vitro* transcription (IVT) on genomic DNA or *in situ* transcription (IST) on fixed cells, with T7 polymerase (*36*, *37*) (**Fig. S1B**).

Once shuffle cassettes are introduced into the genome (*e.g.* randomly via transposition or retrotransposition at a high multiplicity of infection (MOI) or, alternatively, in a targeted fashion), their positions can be efficiently mapped by sequencing T7 IVT-derived transcripts from genomic DNA that contain both cassette-specific barcodes and genomic sequence flanking each integration, with a straightforward protocol that we recently described (*37*) (**Fig. 1D**; **Fig. S1B**). Starting from a parental cell population in which each cell contains a distinct repertoire of integrated shuffle cassettes with mapped genomic locations, Cre recombinase is expected to induce SVs by driving recombination between shuffle cassettes integrated throughout the genome of the same cell. Because these recombination events shuffle which 20 nt barcodes are located *in cis*, specific SVs can be detected and quantified based on novel barcode combinations observed only in “post-shuffle” cells. A key point is that both parental and novel combinations are detectable by simply sequencing PCR amplicons derived from the shuffle cassettes (**Fig. 1E**; **Fig. S1A**). To genotype SVs alongside the capture of single cell transcriptomes, T7 IST can be performed after fixation but prior to scRNA-seq (*36*, *37*), essentially creating an RNA fingerprint of which barcode combinations (and therefore which SVs) are present in association with each single cell transcriptome (**Fig. 1E**). Altogether, this strategy is designed to enable: 1) the multiplex generation of SVs in a population of mammalian cells; 2) the straightforward mapping of the breakpoints and nature of induced SV with no need for whole genome sequencing or karyotyping; and 3) the efficient genotyping and quantitation of SVs, either in bulk (from total DNA or RNA) or at single cell resolution (in conjunction with scRNA-seq).

### *Genome-Shuffle-seq* enables the multiplex generation and haplotype-resolved mapping of thousands of SVs in BL6xCAST mouse ESCs

As a proof-of-concept, we cloned a complex library of shuffle cassettes into a PiggyBac transposon vector (*37*). We then randomly integrated this library into the genome of an F1 hybrid C57BL6/6J × CAST/EiJ (BL6xCAST) male diploid mESC cell line (**Fig. 1C**) (*38*). This cell line was chosen for three reasons. First, a heterozygous variant (SNV or indel) is present, on average, every ∼150 bp, which should facilitate the assignment of shuffle cassette integrations, as well as induced SVs, to one haplotype or the other (*39*). Second, we reasoned that inducing large rearrangements in diploid cells would be less likely to cause cell death, relative to haploid cells. Finally, this mESC line can potentially be differentiated into diverse cell types or organoids, which might eventually facilitate the study of cell type-specific effects of SVs starting from one engineered cell population.

The shuffle cassette library was integrated to BL6xCAST mESC cell line at a high MOI (*40*). After bottlenecking to ∼100 founding clones and re-expanding the cell population, we estimated an average MOI of 123 via quantitative PCR (**Fig. S2A-B**). We identified 9,416 parental barcode combinations in the bottlenecked population by sequencing shuffle cassette-derived PCR amplicons (**Fig. S1A**; **Fig. S2C**). We performed T7 IVT based mapping (*37*) on genomic DNA extracted from this pool to identify the site-of-integration and orientation of each shuffle cassette (**Fig. 2A**; **Fig. S2D**). After filtering out those mapping ambiguously or to multiple locations, we retained 5,088 barcoded shuffle cassettes, well distributed across all chromosomes, whose locations are confidently mapped at base-pair resolution (**Fig. 2A**). ChrX and chrY harbored fewer insertions relative to autosomes, presumably due to their single copy in these male cells and difficulties with mapping on the repetitive chrY (**Fig. 2B**). We used allele-specific SNVs and indels to assign nearly 80% of the shuffle cassettes to either the BL6 or CAST haplotype (**Fig. 2C**; **Fig. S3**). Shuffle cassettes largely mapped to introns and intergenic regions (**Fig. 2D**).

**Fig 2.**
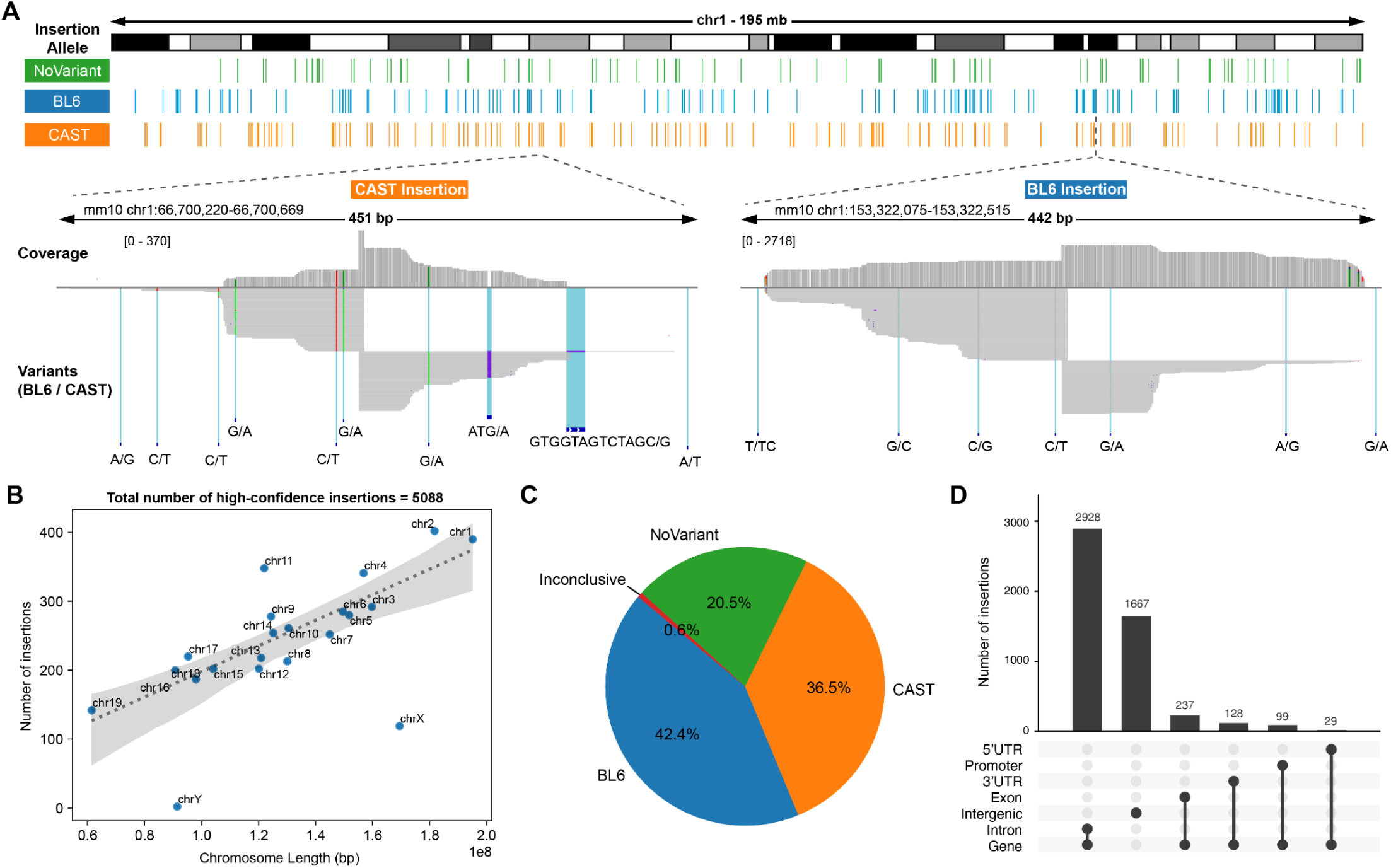
Allele-specific mapping of shuffle cassette insertions. **A)** Insertion sites detected across chromosome 1 in a bottlenecked population of BL6xCAST mESCs, colored by allele. Insets depict pileups of sequencing reads from T7 transcripts for exemplary integrations to the CAST (left) or BL6 (right) haplotype. Alleles are distinguished by the presence of known variants between them. **B)** Number of insertion sites with unique barcodes (*y*-axis) across chromosomes of varying lengths (*x*-axis). The dotted line indicates a linear regression model fit and the shaded gray areas the 95% confidence interval. We have not corrected here for the fact that the X chromosome is single-copy in this male cell line. **C)** Pie chart depicting the distribution of assignments to BL6 or CAST alleles for shuffle cassettes whose genomic coordinates were mapped with high confidence. **D)** UpSet plot of intersection of shuffle cassette integration sites with genomic features.

Next, we next sought to induce SVs, as well as to genotype them via amplicon sequencing of the shuffle cassettes (**Fig. 1**). We transfected varying amounts (200ng, 1μg or 4μg) of either a plasmid expressing Cre recombinase or, as a negative control, the non-targeting Bxb1 recombinase, into cells derived from the bottlenecked population (∼200,000 cells per condition). At 72 hours post-transfection (day 3), cells were harvested and genomic DNA isolated. We then PCR amplified and deeply sequenced shuffle cassette barcode pairs, and assessed whether new combinations were present (**Fig. 3A**). As we hoped, while almost no non-parental barcode combinations were detected in the non-targeting Bxb1 recombinase control condition, >5,000 novel barcode combinations were detected across the conditions involving Cre recombinase (**Fig. 3B**). Of note, because in some cases we detect both non-parental barcode combinations generated by the recombination event (**Fig. 1E, Fig. S1C-D**), these correspond to 4,856 inferred, unique SV events.

**Fig 3.**
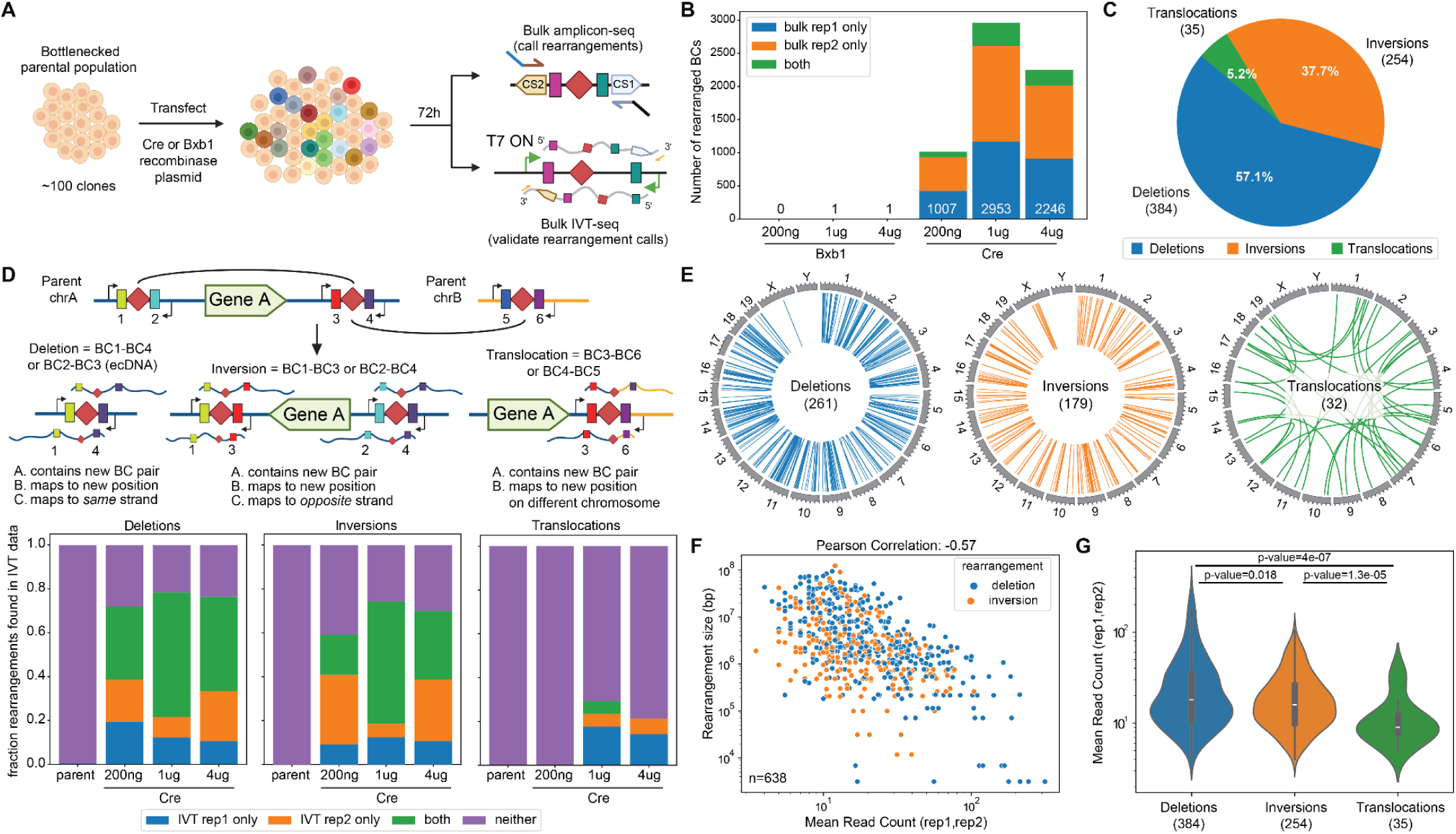
Multiplex induction and efficient genotyping of large-scale rearrangements throughout a mammalian genome. **A)** Experimental schematic. **B)** Number of novel (*i.e.* non-parental) barcode combinations at >1 UMI detected in each condition from amplicon-seq data. Different colors indicate those rearranged barcode combinations found in one technical replicate or both within each condition. **C)** Pie chart showing the distribution of SV types that are detected in both technical replicates of a Cre transfection sample. SVs detected in multiple conditions are counted independently. **D)** Schematic of approach for validation of SV calls using matched IVT-seq data from the same sample (top). The proportion of each SV type that is supported by at least one read in the IVT-seq data are depicted below. **E)** Circos plots of the unique set of SVs that are shared between technical replicates of each sample. SVs detected in multiple conditions are counted once. **F)** Scatter plot of rearrangement size (*y-*axis) vs. mean read count (*x-*axis) for deletions and inversions detected at day 3. Pearson correlation is calculated between the log10 values of the two metrics. **G)** Violin plots depicting the distribution of read counts for deletions, inversions and translocations detected at day 3. Inset within each violin plot is a box plot of the distribution with the median value depicted as a white line, the length of the box depicting the interquartile range and the whiskers depicting the extent of the distribution. P-values are calculated using the non-parametric Mann-Whitney U test.

We are likely only observing a small fraction of the SVs that could potentially be generated by this approach. First, nearly all (∼99.9%) of amplicons in conditions involving Cre recombinase matched “parental” barcode combinations, consistent with SVs being individually and collectively rare in the cell population (**Fig. S4A**). Second, the vast majority of novel barcode combinations were not shared between technical replicates prepared from different genomic DNA aliquots from the same Cre condition, nor across Cre conditions. Thus, we likely would have detected many more SVs simply by processing more Cre-exposed cells from this same population of ∼100 founding clones (each bearing ∼50 confidentally mapped and oriented shuffle cassette integrations).

### *Genome-shuffle-seq* induces thousands of unique deletions, inversions and translocations

For each novel barcode combination, we can infer the class and size of the corresponding SV based on the relative genomic coordinates and orientation of the parental shuffle cassettes (**Fig. 1E**; **Fig. S1C-D**). For the subset of SVs shared by both technical replicates of a given Cre transfection condition, 53% were observed in at least one other transfection condition (**Fig. S4B**). If we focus on SVs observed in both technical replicates of a condition (n=673), deletions and inversions are much more common than translocations (**Fig. 3C**). However, if we consider all detected SVs (n=6879), translocations comprised the majority (**Fig. S4C**). We return to the potential interpretations of this difference further below.

SVs involving all chromosomes were detected, with the exception of chrY (**Fig. 3E**; **Figs. S5-S6**). As expected, the number of SVs detected per chromosome was correlated with chromosome size, presumably a consequence of the number of shuffle cassette insertions (**Fig. S6A-B**). Interestingly however, when we break this down by SV class, some chromosomes appear enriched or depleted for certain types of rearrangement (**Fig. S6C-H**).

For deletions and inversions, there was an exponentially inverse correlation between SV size and its abundance, as inferred by the number of sequence reads supporting the corresponding barcode combination (**Fig. 3F**; **Fig. S4D**). The subset of deletion/inversion SVs supported by both technical replicates (n=638) had a read-counted weighted median event size of ∼1 Mb, while the complete set (n=3163) had a larger median event size ∼2.5 Mb (**Fig. S4E**). This may reflect the known property of Cre recombination efficiency to drop exponentially with genomic distance in mammalian cells (*25*) and/or selection acting against those cells containing large genomic deletions or inversions.

To orthogonally validate the SVs inferred from novel barcode combinations, we performed IVT-seq (*37*), *i.e.* the same protocol with which the coordinates of the parental shuffle cassettes were originally mapped, on “post-rearrangement” genomic DNA. Given the inward orientation of the T7 sites, we expect IVT transcripts from each T7 site to cover not only the novel barcode combination, but also flanking genomic DNA, thereby facilitating direct verification of the corresponding SV (**Fig. 1B**). For deletion/inversion SVs, the majority of either the technically replicating (n=638) or full (n=3163) sets of deletions were validated by at least one IVT-seq read from the same condition, but entirely unsupported by IVT-seq data from parental cells (**Fig. 3D**; **Fig. S4F**). In contrast, despite the fact that translocations made up the majority of all detected SVs, a much smaller fraction of translocations were supported by IVT-seq data (**Fig. 3D**; **Fig. S4F**). Consistent with that, in the amplicon sequencing of shuffle cassettes that originally detected each SV, translocations were supported by substantially fewer reads than deletions or inversions (**Fig. 3G**; **Fig. S4G**). Artifactual explanations such as chimeric PCR are ruled out by the essentially complete absence of reads supporting any type of SV, including translocations, in Bxb1-treated control cells (**Fig. 3B**). One potential explanation of the both lower abundances and lower validation rates for translocations is that many detected deletions and inversions are being generated recurrently even within a single condition/replicate (*i.e.* in independent cells), while detected translocations occur uniquely, precluding validation in independent aliquots of “post-rearrangement” genomic DNA and lowering read counts within the aliquot in which they occurred. An alternative explanation is that translocations are occurring at similar rates but being strongly selected against, either indirectly (via generalized Cre toxicity) or directly (via phenotypic consequences of the translocation itself).

Taken together, these results show that we are able to induce, detect, quantify and characterize thousands of deletions, inversions and translocations in a pool of cells in a single multiplex experiment with *Genome-Shuffle-seq*, without any single-cell cloning, genotyping or whole genome sequencing. However, the basis for the differential recurrence rates, validation rates, and supporting read depths for various SV classes remained unclear.

### Cre-mediated rearrangements are rapidly depleted from the *in vitro* mESC culture

To further investigate this, we sampled the population of Cre-transfected cells at days 5 and 7 post-Cre transfection, and sequenced shuffle cassette-derived amplicons to analyze the fate of cells induced to generate SVs on day 3 (**Fig. S7**; **Methods**). We observed a marked decline in the number of detected SVs at later time points, with almost no SVs detected by day 7 (**Fig. S7C-D**). We first hypothesized that this could be due to the well-documented toxicity of Cre recombinase to mammalian cells, which is thought to impose cost on cell fitness in proportion to the number of target sites in the genome, in a p53-dependent manner (*41–43*). The impact of this generalized Cre toxicity on transfected cells would lead to untransfected or poorly transfected cells overtaking the population. As a potential solution, we hypothesized that tamoxifen-inducible Cre variants (CreERT2 and ERT2CreERT2) could be used to restrict the time window of Cre activity, limiting toxicity (*41*, *44*). To test this, we transfected bottlenecked parental cells with inducible Cre variants, treated with 0.5 μM tamoxifen for 24 hours at 1 day post transfection, and collected samples at days 3, 5 and 7 and performed amplicon sequencing of shuffle cassette barcodes (**Fig. S7A**). Both inducible Cre variants induced far fewer SVs than constitutive Cre, and also failed to facilitate survival of cells bearing SVs at day 7 (**Fig. S7C-D**). As an alternative strategy, we sought to reduce cell death by treating cells with the p53 inhibitor Pifithrin-α (20μM) for 48 hours post-Cre transfection. We chose not to prolong p53 inhibition beyond that due its toxicity and effects on stem cell maintenance and differentiation (*45*, *46*). Although Pifithrin-α treatment increased the number of detected SVs at day 3, we once again observed a marked decline in the abundance of SVs by day 5 (**Fig. S7E-F**).

Another potential explanation for the decrease in SV abundance over time is that the Cre-induced SVs are themselves causing fitness defects, such that cells with less phenotypically consequential SVs, or those lacking SVs entirely, outcompete them in the population. If this were the case, sorting out rearranged cells into single wells such that they can grow out in isolation would be expected to solve this problem. We co-transfected either Cre or Bxb1 recombinase and a Cre-reporter that conditionally expresses the red fluorescent protein (RFP) into the bottlenecked parental population. Transfected cells were either treated with Pifithrin-α or no drug for 48 hours post-transfection. On day 3, 720 RFP-positive cells from the Cre condition samples were sorted into single wells of 96 well plates that either contained Pifithrin-α or no drug (**Fig. S8A-C**). Pifithrin-α treatment led to a marked increase in the number of clones that grew up after single-cell sorting from the Cre-transfected population, consistent with p53 inhibition reducing cell death (**Fig. S8D**). However, no SV-supporting barcode combinations were detected upon sequencing barcode pairs in genomic DNA derived from 86 single cell clones. Interestingly, the median number of parental shuffle cassettes detected per Cre-treated sample was lower than for Bxb1 samples (**Fig. S8E**). This suggests that clones with higher numbers of integrated shuffle cassettes may be selected against after Cre transfection.

### Genotyping induced SVs at single cell resolution in scRNA-seq data

*Genome-shuffle-seq* was intentionally designed for compatibility with the efficient genotyping of SVs in conjunction with single cell transcriptomes (**Fig. 1E**). To test this aspect of our scheme, we sorted a pool of Cre-transfected cells at 72 hours post-transfection for cells positive for Cre reporter activity, and then subjected these to fixation, IST and scRNA-seq. For this experiment, we combined cells treated with vs. without Pifithrin-α, and also included an independent sample from parental cells as a control (**Fig. 4A**). Sorted cells were fixed with methanol followed by T7 IST (*36*, *37*). At this stage, fixed cells are expected to contain both endogenous mRNAs and T7-derived transcripts that cover the shuffle cassette barcode pairs. To capture both sets of transcripts with a common cell barcode (cell BC), we subjected the cells to 3’ scRNA-seq with feature barcoding on the 10X Genomics platform (**Fig. S9A**).

**Fig 4.**
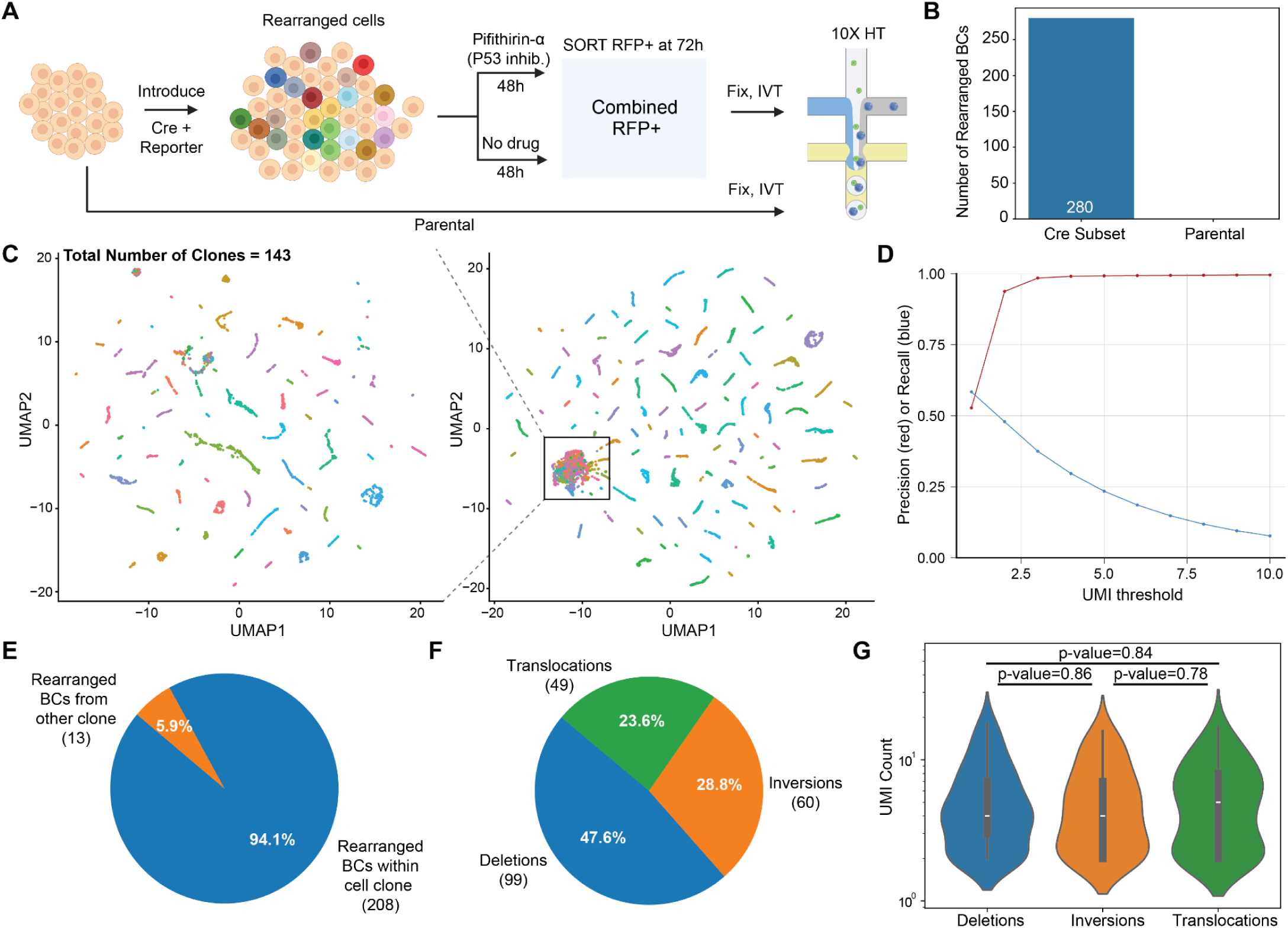
Detection of rearrangements in single cells. **A)** Experimental schematic. At 72h post transfection, cells were sorted based on the activity of the Cre reporter, methanol-fixed, subjected to T7 IVT and then sc-RNA-seq on the 10x Genomics platform. **B)** Number of novel barcode combinations detected either in the whole parental sample or a downsampled subset of the Cre sample in sc-RNA-seq. **C)** 9412 cells were successfully assigned to 143 clonotypes. Here we visualize 8522 of these assigned cells with at least 20 T7 UMIs in UMAP space based on the T7-derived barcodes that are detected at >1 UMI, colored by clonotype assignment. Plot on the left was generated by iterative clustering of subset of cells from global UMAP shown on right. **D)** Mean precision and recall of detected T7 barcodes in cells compared to the expected barcodes list from respective clonotypes to which the cells are assigned (*y-*axis), at various UMI thresholds (*x-*axis). **E)** Pie chart depicting the fraction of rearranged barcodes (BCs) detected in cells that are congruent with the clonotype assignment of that cell. **F)** Pie chart depicting the distribution of rearrangement types detected in single cell data. G) Violin plots depicting the distribution of UMI counts per rearrangement type in single cell data. The inset is a box plot of the distribution with the median value depicted as a white line, the length of the box depicting the interquartile range and the whiskers depicting the extent of the distribution. P-values are calculated using the non-parametric Mann-Whitney U test.

We were able to recover high-quality transcriptomes from our Cre-treated sample, indicating that fixation and IVT upstream of droplet formation did not compromise the remainder of the protocol. After filtering cells by mitochondrial content and the number of transcriptome UMIs detected, we recovered ∼15,000 and ∼19,000 scRNA-seq profiles from Cre-treated and parental samples, respectively, although the parental scRNA-seq data was lower quality due to a wetting failure during droplet generation (**Fig. S9B**). In the Cre-treated sample, we detected a median of 108 T7 UMIs per cell BC, which reflected a median of 46 unique shuffle barcode combinations per cell (**Fig. S9C-D**). When restricted to those barcode combinations that could be confidently mapped to a unique genomic location, we recovered a median of 22 barcode combinations per cell.

To assess rearrangements, we compared the shuffle cassette barcode combinations observed in T7 IST + scRNA-seq data to the parental barcode pairs, which highlighted 1123 novel barcode combinations among ∼15,000 scRNA-seq profiles (**Fig. S10A**). We confirmed that these novel barcode combinations were not an artifact of scRNA-seq library construction by downsampling scRNA-seq data from the Cre-treated condition and comparing to scRNA-seq data from the parental condition. At similar depths, 280 novel barcode combinations were observed in the Cre-treated condition, while none were observed in the parental condition (**Fig. 4B**).

Given the fact that we permeabilize cells for T7 IST and the short length of T7-derived transcripts, our protocol may be at heightened risk for ambient RNA contamination, an established confounder of scRNA-seq (*47*, *48*). To investigate this, we first identified a set of 143 “clonotypes” through iterative clustering on the combinations of barcodes observed in single cells, leveraging both scRNA-seq data as well as amplicon-seq data from single cell clones (**Fig. S8**). These clonotypes essentially correspond to the clones defined by the bottlenecked parental population (**Fig. 3A**), each of which is expected to be defined by a unique complement of PiggyBac integrations, and therefore a unique combination of barcodes.

We then sought to assign each cell to a clonotype based on the set of T7 IST-derived barcodes detected in that cell. Cells were considered successfully assigned if >10% of the barcodes from the clonotype were recalled in that cell and >75% of the total barcodes detected in that cell were precisely from that clonotype. We further filtered cells whose second-best clone assignment had >10% recall. This analysis was designed to remove cells with low barcode capture, doublets and those belonging to undetected clonotypes, leaving a set of 9412 cells that could be confidently assigned to one of the 143 clonotypes (**Fig. S10B**). This subset of cells clustered neatly in UMAP space based solely on their complement of T7 IST-derived barcode combinations (**Fig. 4C**).

Within the set of clone-assigned cells, we estimated a UMI cutoff for confident barcode detection by performing a precision-recall analysis at various UMI thresholds. We observed a large jump in the precision of barcodes detected in a cell when compared to the clonotype barcodes at a UMI threshold of >1 without much cost in terms of recall loss (**Fig. 4D**). Interestingly, although not obviously different from other cells based on their transcriptome quality (**Fig. S9B**), the clonotype assignment rate of cells with rearranged barcode combinations at >1 UMI (n=320) was modestly lower than that of clonotype-assigned cells lacking evidence for any SV (**Fig. S10B**).

Finally, we asked whether the novel barcode combinations detected in a cell were congruent with the clonotype identity assigned to that cell. Indeed, ∼94% of the 221 rearranged barcodes detected at >1 UMI in clone-assigned cells comprise a novel barcode combination involving parental barcodes from the same clonotype (**Fig. 4E**). In order to be conservative, we proceeded with 208 novel barcode combinations that were: 1) detected in cells that were confidently assigned to a clonotype, 2) composed of individual barcodes present in the original clonotype; and 3) detected at >1 UMI in that cell (**Fig. S10A-B**). Collectively, these data suggest that we can detect, map and confidently assign induced SVs to single cells with associated transcriptomes.

Similar to our analysis shown in **Fig. 3**, we were able to infer the nature of each of the 208 SVs based on the parental locations of the barcodes contributing to each novel pair, which once again included deletions, inversions and translocations (**Fig. 4F**; **Supplementary Movie 1**). Most SVs were only detected in only one cell (range 1-4) and most cells only contained a single detected SV (range 1-7) (**Fig. S10C-D**). UMI counts were not different between rearrangements of each class within these single cells (**Fig. 4G**). This contrasts with the lower read counts observed for translocations in bulk data (**Fig. 3G**), strongly suggesting that these were driven by cellular abundance differences that do not manifest at single cell resolution.

### Hundreds of ecDNAs are launched and readily detected by *Genome-Shuffle-seq*

Cre-mediated deletions between loxPsym sites *in cis* result in the formation of a single genomic scar which reflects a deletion of the intervening sequence and a single extra-chromosomal DNA circle (ecDNA) that contains the intervening sequence (**Fig. 5A**). Both species are expected to be present in equal stoichiometry at the time of their formation. We were able to assign each novel, deletion-indicative barcode combination detected by either sequencing of PCR amplicons or in scRNA-seq data as deriving from the scarred chromosome or ecDNA circle, based on the relative orientation of the parental shuffle cassette insertions and the pair of rearranged barcodes that was detected (**Fig. 1E**). To our surprise, barcodes derived from the ecDNA species, rather than genomic deletion scars, comprised the majority of deletion detection events in PCR amplicon data, an asymmetry that was even more exaggerated in the scRNA-seq data (**Fig. 5B**). Further, for deletions for which we were able to detect both the ecDNA and deletion scar in the same sample in PCR amplicon data, ecDNAs were detected at 2-3-fold higher read counts (**Fig. 5C**). Only 3 shared ecDNA-genomic pairs were detected in the same cell in our single-cell data, precluding a similar analysis. Additionally, detected ecDNAs tended to be larger than detected deletion scars (**Fig. 5D-E**; **Fig. S11**).

**Fig 5.**
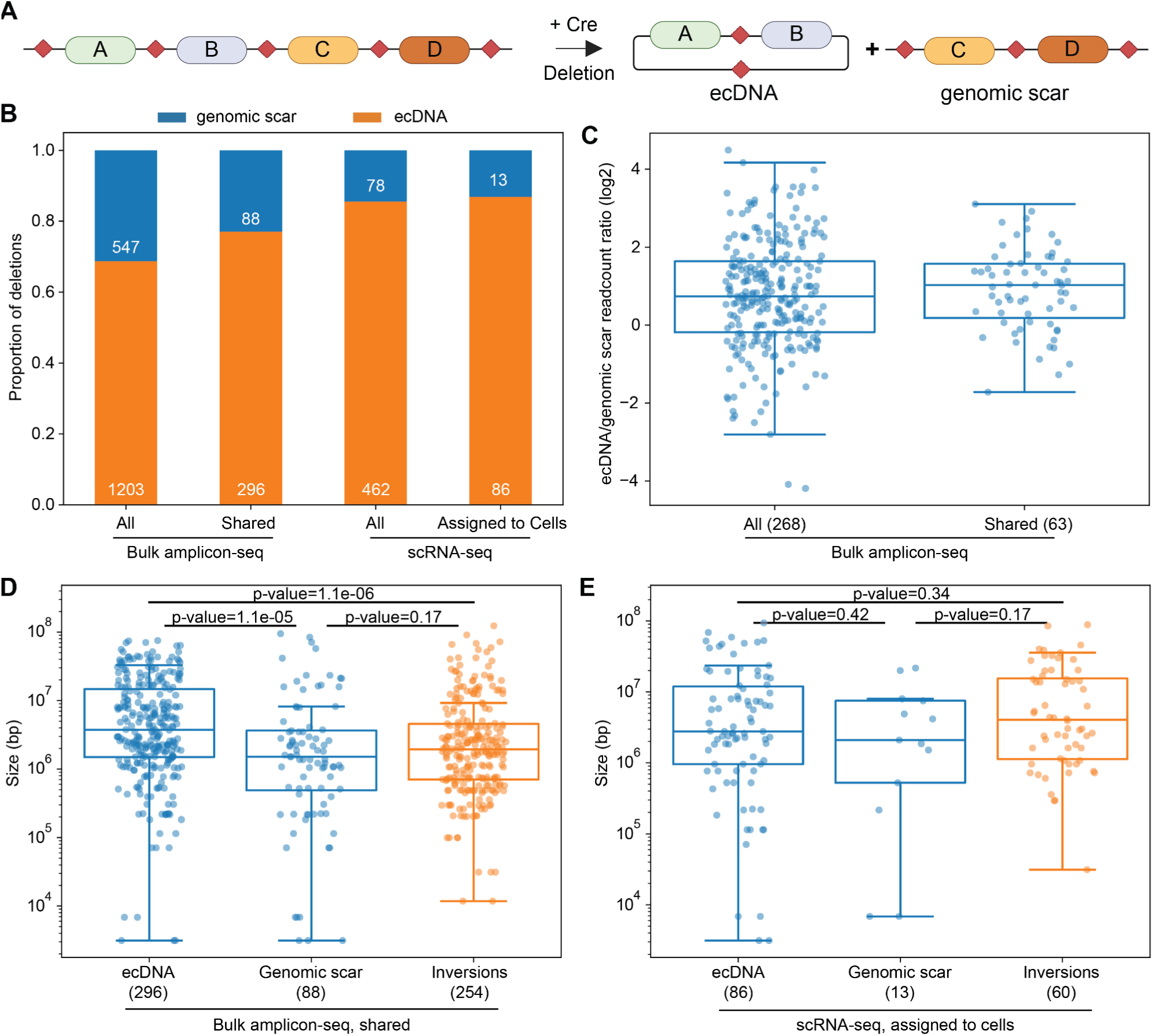
Hundreds of ecDNAs are launched by *Genome-shuffle-seq*. **A)** Schematic of ecDNA formation during Cre-mediated deletion. The genomic deletion and ecDNA species are expected to have 1:1 stoichiometry at the time of formation. **B)** Barplot depicting the proportion of rearrangements called as deletions that are represented by the set of barcodes expected to be on the ecDNA vs the genomic deletion scar. Deletions from bulk amplicon-seq at 72h post Cre transfection (all deletions and those present in both technical replicates) and those detected in scRNA-seq (all detected deletions and those assigned to cells) are included. **C)** Boxplot of the log2 ratio of read counts for the barcode pairs that represent the ecDNA vs. genomic copy for the set of ecDNA-genomic deletion pairs that were both detected in the same bulk amplicon-seq sample at 72h (from all deletions and those shared between technical replicates). The horizontal solid line indicates the median, the length of the box depicts the interquartile range of the distribution and the whiskers depict the rest of the distribution excluding outliers. **D)** and **E)** Size of each ecDNA, genomic copy and inversion detected in bulk amplicon-seq (shared between technical replicates) and scRNA-seq (assigned to cells) data. Boxplot elements are the same as in panel **C**. Depicted p-value is calculated using the non-parametric Mann-Whitney U test.

This unexpected deviation from the expected 1:1 stoichiometry at the time of rearrangement formation has a number of potential explanations. First, ecDNAs may be preferentially isolated and purified compared to the native chromosomes owing to their smaller size and circular topology in bulk genomic DNA preparations used as templates for PCR. Second, in the case of our scRNA-seq data, SV detection is dependent on T7 IST, which may preferentially transcribe from the ecDNA owing to more highly accessible chromatin states (*49*). Third, it is possible that ecDNAs launched from the genome undergo amplification through either asymmetric segregation or uninhibited replication as compared to native chromosomes, which in combination with selective pressures to maintain copy number could contribute to the observed asymmetry (*50*, *51*). Further work is necessary to distinguish between these technical vs. biological explanations.

## Discussion

In this study, we describe *Genome-Shuffle-seq,* a novel method for the inducible generation and facile characterization and quantitation of thousands of mammalian SVs within a pool of cells, without any need for clonal isolation and genotyping nor for whole genome sequencing. We also show how SV identities can be captured alongside single-cell transcriptomes for hundreds of such rearrangements. Finally, we show that *Genome-Shuffle-seq* can be used to produce synthetic ecDNAs in mammalian cells, and that these species are also detectable in scRNA-seq data.

As currently described, *Genome-Shuffle-seq* has several key limitations. First, we are unable to detect the ∼50% of rearrangements that result in the formation of a shuffle cassette with the same capture sequence on either side of the loxPsym site, presumably due to suppressive PCR (**Fig. S1C-D**) (*52*, *53*). Alternative cassette designs that are longer and include more sequence diversity or employ PCR-free methods for library generation may help address this issue. Second, the capture rates of T7-derived transcripts in scRNA-seq assay were limited; higher rates would facilitate more comprehensive and higher confidence detection of induced SVs, and ideally the complete *in silico* karyotype of each profiled cell. Capture rate improvements are expected with an updated protocol in which construct-specific primers are spiked in during the initial cDNA library amplification (*54*, *55*).

A third limitation relates to the rapid depletion of SVs within one week of their induction. We had sought to obtain clones derived from cells with a given SVs (even within a polyclonal population) to facilitate the robust detection of gene expression changes in scRNA-seq data, *i.e.* to overcome the the dropout rates and sparsity inherent to this type of analysis, and assign rearrangements to changes in gene expression with some measure of statistical confidence (54). However, in scRNA-seq data, cells containing rearrangements were rare, and the vast majority of rearrangements were only detected once, consistent with the depletions of SVs detected in bulk.

Stepping back, the landscape of observed SVs, including their relative abundances as a function of time, are likely shaped by three forces: 1) the efficiency of SV formation; 2) selection acting upon induced SVs; and 3) generalized Cre-mediated toxicity. In our view, the most parsimonious explanation of the patterns observed in our data involves a combination of explanations (1) and (3). In particular, we hypothesize that the varying abundances of SV classes are primarily due to differential rates of recurrence. Intrachromosomal deletions and inversions are both abundant and recurrent within and across conditions, in a manner inversely correlated with size, due to the well-established properties of SSRs, which favor events between closely located recognition sites *in cis* (*25*). On the other hand, among all unique SVs detected, translocations are most common, because there is a combinatorially larger number of possible translocations than inversions or deletions. Translocations are supported by fewer reads and do not replicate, because that same translocation, involving the same pair of shuffle cassettes, is extremely unlikely to have recurred, either within or across samples.

The well-documented toxicity of Cre, rather than selection against larger intrachromosomal events or translocations, seems to be the simplest explanation for the rapid depletion of SVs within a week of their induction. The alternative explanation, selection against the SV events themselves, seems less likely, given the diploid nature of the cell line, the results of our single cell cloning experiments, and the fact that even small SVs were not detectable one week after Cre induction. We are using p53-competent stem cells, which may be particularly sensitive to Cre toxicity. Our cells also harbor a large number of SSR recognition sites, which may further increase their sensitivity. Looking forward, we anticipate several approaches that might be taken to reduce Cre toxicity and/or boost recovery of successfully shuffled genomes, including: 1) switching to a p53-null cell line; 2) tighter regulation of recombinase expression; 3) switching to a recombinase with lower fitness costs than Cre (*56*, *57*); 4) including a conditional selection marker that would be reconstituted upon shuffle cassette recombination (*24*, *25*); 5) lowering the number of SSR recognition sites per cell; and/or 6) lowering the distances between SSR recognition sites.

Despite these remaining challenges, we believe that *Genome-Shuffle-seq* lays the foundation for large-scale single-cell genotype-to-phenotype screens for the impact of thousands to millions of mammalian SVs and ecDNA species on gene expression, chromatin structure and genome organization, analogous to Perturb-seq or CROP-seq (*58*, *59*). As a related approach, the targeted introduction of shuffle cassettes to individual mammalian genomic loci, for instance by bottom-up assembly (*60*), would facilitate the dissection of regulatory element interactions and locus architecture in specifying gene regulation. *Genome-shuffle-seq* could also readily be adapted to study the cell-type specific impact of SVs by differentiating a single engineered population into *in vitro* multi-cellular models or *in vivo* using whole organism models. Of note, in related work conducted independently, Koeppel, Ferreira and colleagues describe a complementary strategy for the “randomization” of mammalian genomes with engineered SVs using highly multiplexed prime editing-mediated insertion of loxPsym sites (*61*). Beyond enabling the more systematic study of SVs and ecDNAs, these approaches may also serve as an entry point for the engineering of a ‘minimal genome’ comprising the essential complement of genetic information required for propagation of any mammalian cell, potentially useful as a universal chassis for cell-based therapy (*62*).

## Supplementary Figures/Movie

**Fig S1.**
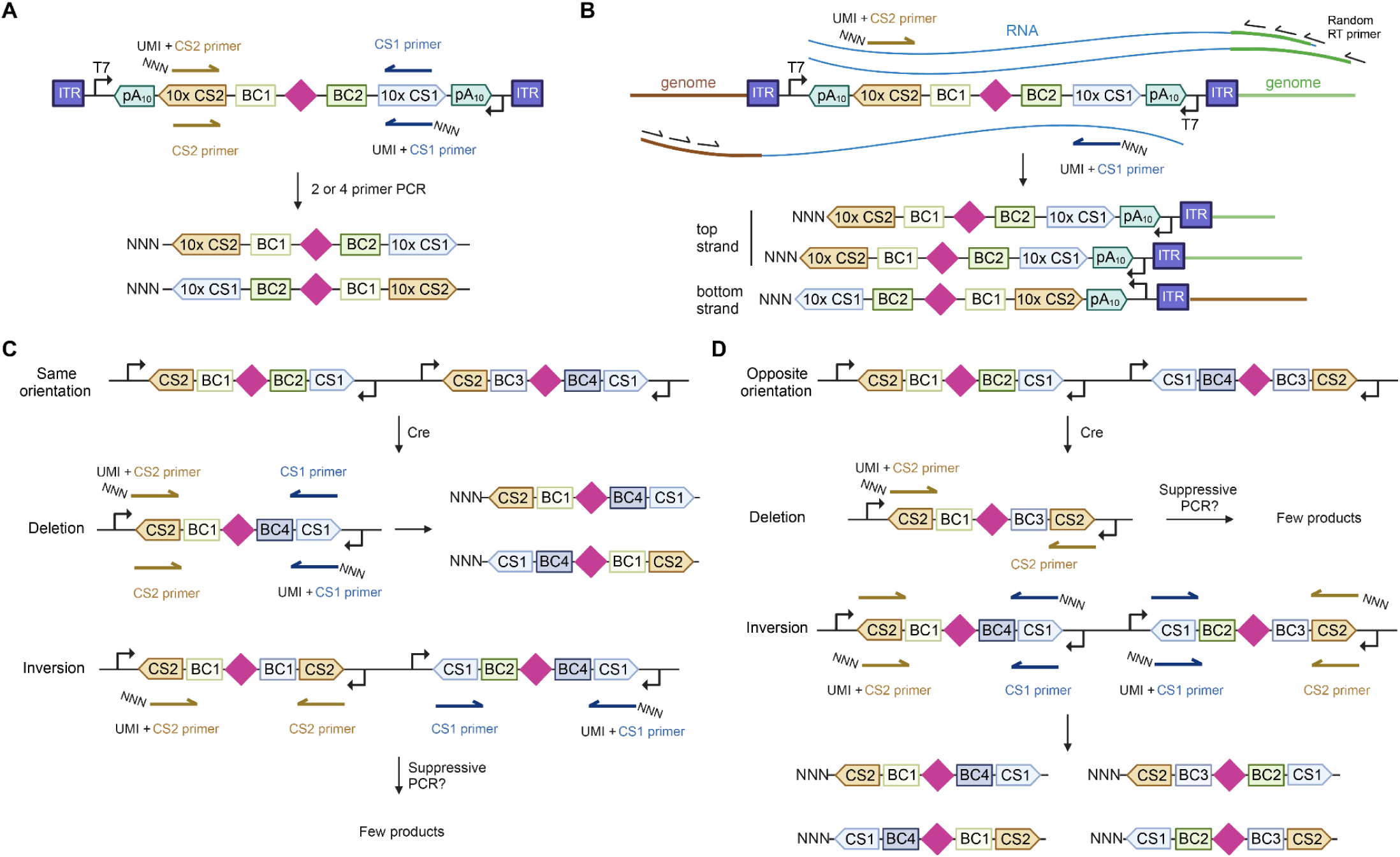
Schematic of sequencing library construction strategies for amplicon-seq and IVT-seq. **A)** Schematic of the shuffle cassette bearing the loxPsym site (diamond) flanked by two barcodes (BC1, BC2). Two 10x Genomics capture sequences (CS1, CS2) serve as binding sites for PCR primers for sequencing library construction. Only amplicons generated using one unique molecular identifier (UMI)-containing primer and one non-UMI primer can cluster and be successfully sequenced on an Illumina flow cell due to the sequencing adapters they encode. **B)** After IVT, transcripts are generated from both the top and bottom strand T7 promoters, and are expected to contain both BC1 and BC2 as well as adjacent genomic sequences from one side of the integrated shuffle cassette. Reverse transcription (RT) is performed with a primer containing 8 random bases at its 3’ end. PCR is performed with a UMI-containing primer and one primer annealing to the constant sequence in the RT primer to yield the final sequencing library. **C)** and **D)** Recombination between two insertions *in cis* can lead to an inversion or deletion with shuffle cassettes containing either the same or different capture sequences. Theoretically, PCR products from a cassette with the same CS should amplify and cluster on an Illumina flowcell when libraries are generated using all 4 primers. However, empirically we find that these products are not readily detected, probably due to suppressive PCR (*52*, *53*) (see **Fig. S7B**).

**Fig S2.**
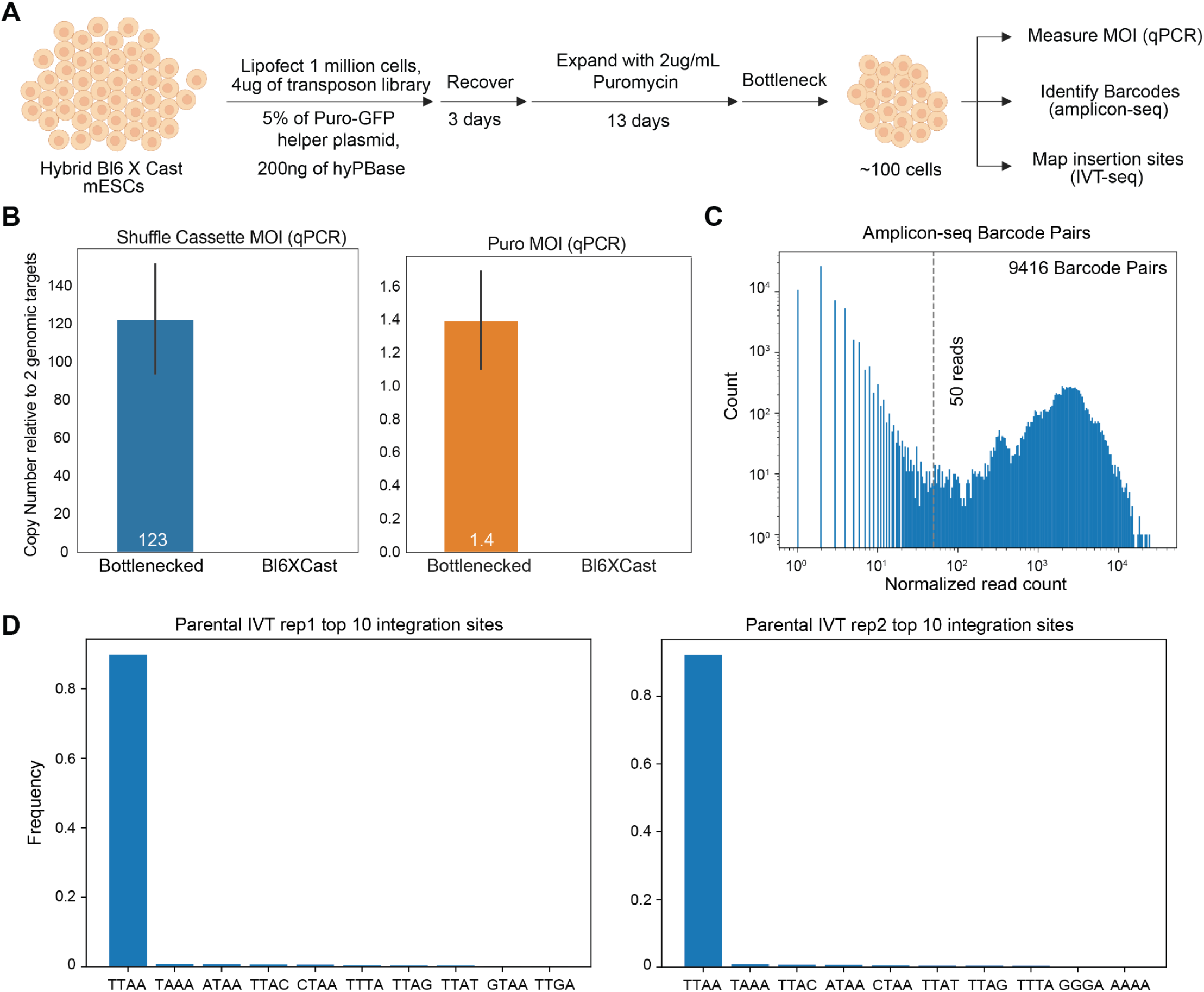
Integration and characterization of shuffle cassette library into mESCs. **A)** Schematic of experiment to integrate shuffle cassettes to the genomes of mESCs at a high multiplicity of infection (MOI) by co-transfection with a small percentage of a helper plasmid containing the puromycin resistance gene (*40*). **B)** Copy number of shuffle cassettes and the puromycin resistance gene were estimated in the bottlenecked population via quantitative PCR (qPCR) relative to two genomic targets (Trfc, Tert). The height of the bar represents the mean and the error bars indicate the standard deviation of the copy number measured relative to the two genomic targets. **C)** Histogram of read count for each barcode pair detected in amplicon-seq data normalized to sequencing depth across 4 technical replicates. **D)** Frequency of the first 4 bp of the genomic sequence detected in IVT-seq reads in technical replicate 1 and 2 from parental cells. TTAA is the expected sequence given our use of the PiggyBac transposon.

**Fig S3.**
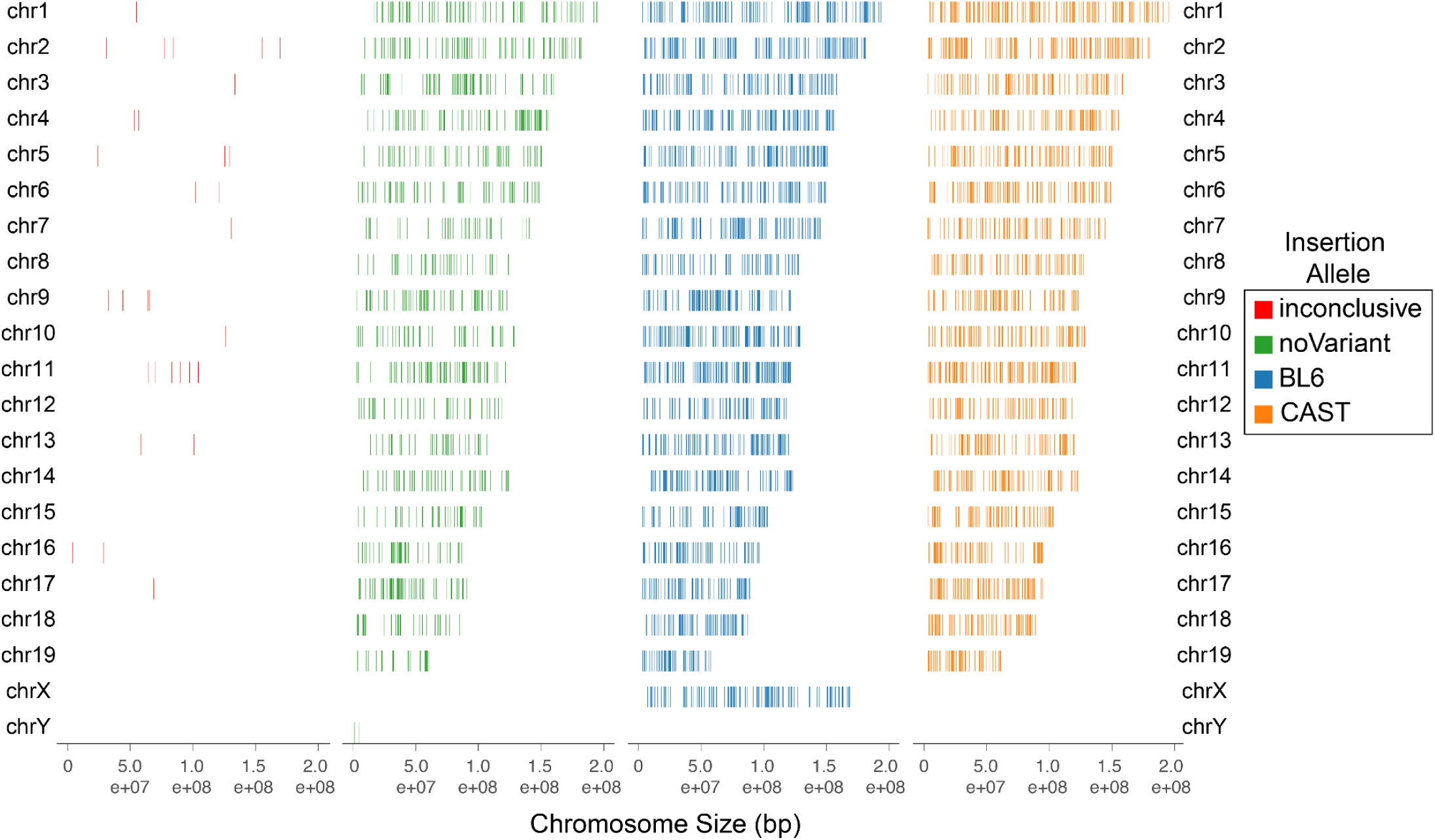
Allele-specific insertion sites across all chromosomes. Insertion sites across all chromosomes for shuffle cassettes whose genomic coordinates were mapped with high confidence, colored by allele. Inconclusive indicates that there is conflicting evidence for the insertion allele, while noVariant denotes those insertions that were un-assigned due to a lack of reads that overlap with a known variant between the BL6 and CAST genomes.

**Fig S4.**
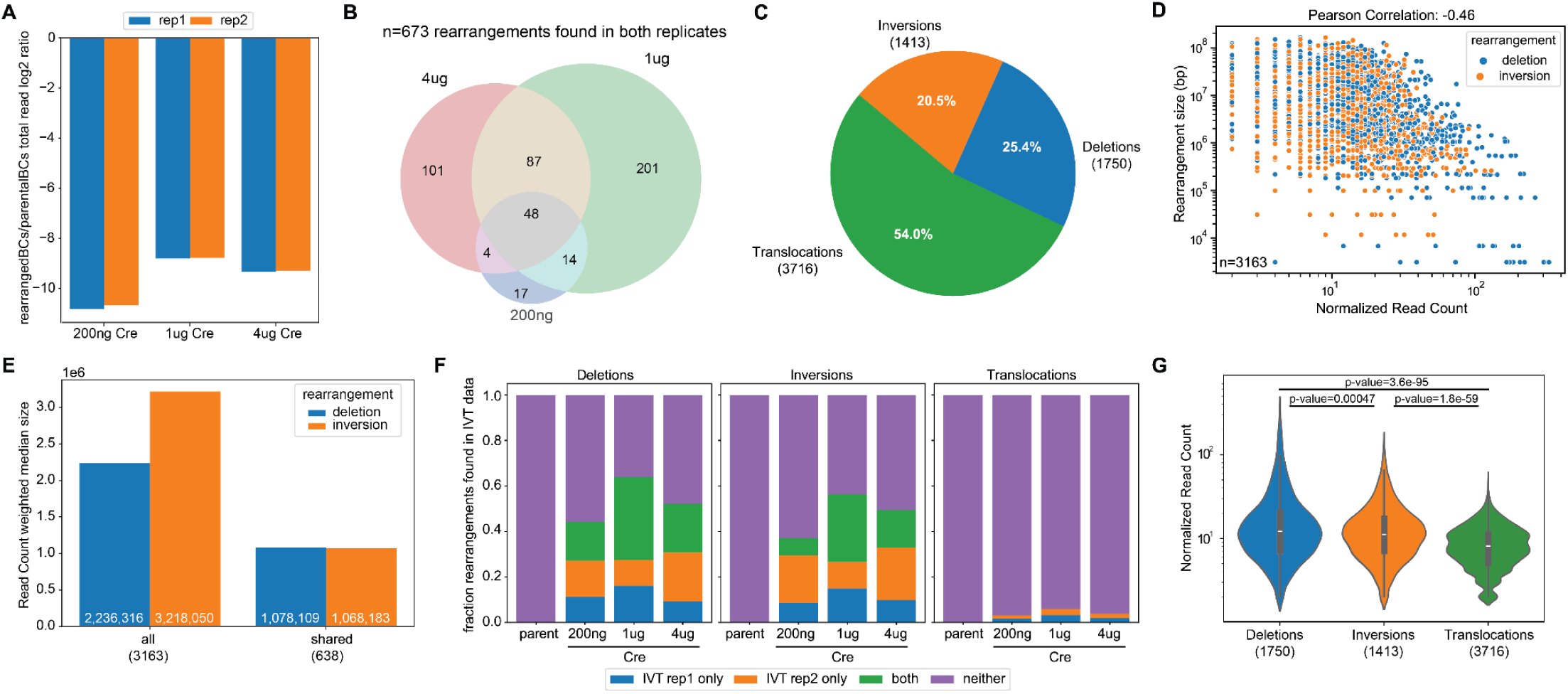
Characteristics of the complete set of rearrangements detected in bulk by amplicon-seq at 72h post-Cre transfection. **A)** Log2 ratio of total reads that contain rearranged barcode (BC) pairs to the total number of reads that contain parental BC pairs in technical replicates of each Cre transfection sample. **B)** Venn diagram depicting the overlapping relationships between Cre transfection samples for the subset of SVs that are detected in both technical replicates of each sample. **C)** Pie chart depicting the distribution of SV type for all rearrangements detected at 72h. **D)** Scatter plot of rearrangement size (*y-*axis) vs. normalized read count (*x-*axis) for deletions and inversions detected at day 3. Pearson correlation is calculated between the log10 values of the two metrics. **E)** Median size of inversions and deletions, weighted by their read count, for both the complete set of rearrangements (left) and those shared between technical replicates for a condition (right). **F)** Similar to lower part of Fig. 3D, the bar plot shows the proportion of each SV type (from the complete set of rearrangements at 72h) that is supported by at least one read in the IVT-seq data. **G)** Violin plots depicting the distribution of read counts for deletions, inversions and translocations for the complete set of rearrangements detected at day 3. Inset within each violin plot is a box plot of the distribution with the median value depicted as a white line, the length of the box depicting the interquartile range and the whiskers depicting the extent of the distribution. P-values are calculated using the non-parametric Mann-Whitney U test.

**Fig S5.**
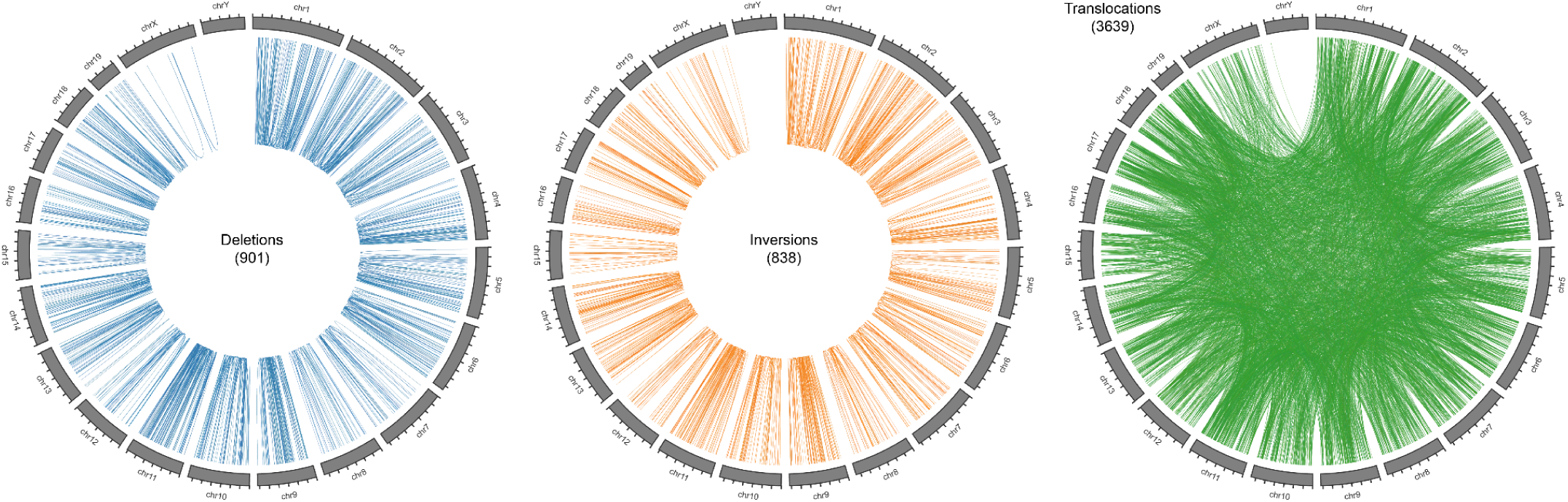
Circos plots of all rearrangements detected at 72h post-Cre transfection. Depicted rearrangements are from across all samples, including those that are not shared between technical replicates.

**Fig S6.**
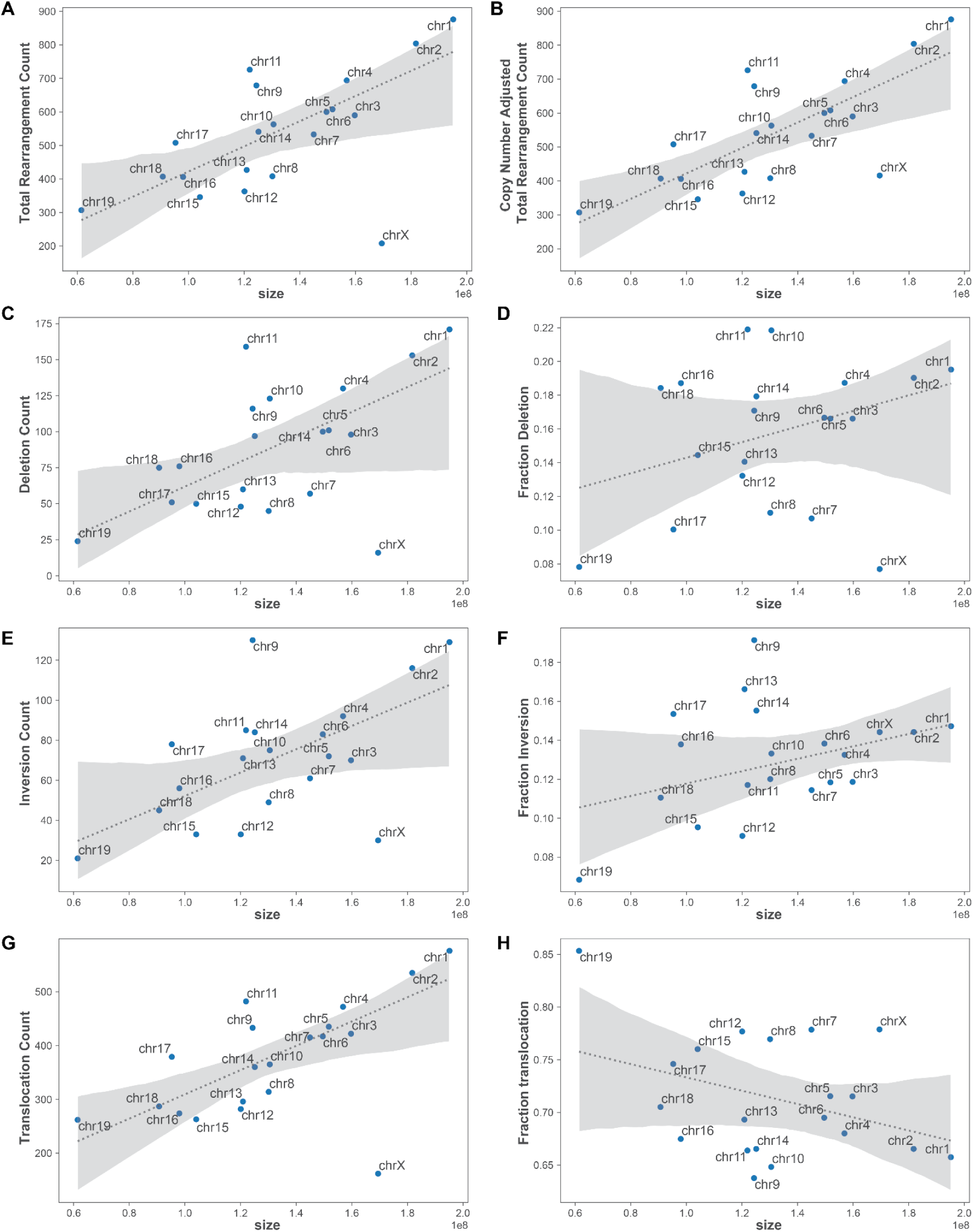
Distribution of number or fraction of rearrangements of each type across chromosomes. All plots in this figure depict a scatter plot of the distribution of all rearrangements detected at 72h post-Cre transfection across mouse chromosomes except chrY, with the size of the chromosomes depicted along the *x-*axis. The dotted line indicates linear regression model fit and the shaded gray areas the 95% confidence interval. The *y*-axis of each panel is either: **A)** the total number of events detected; **B)** the total number of events detected with the number of events on the X chromosome multiplied by 2 to normalize for copy number; **C)**, **E)**, **G)** the total number of deletions **(C)**, inversions **(E)** or translocations **(G)**, respectively; **D)**, **F)**, **H)** the proportion of rearrangements on a given chromosome that are deletions **(D)**, inversions **(E)** and **(G)** translocations, respectively.

**Fig S7.**
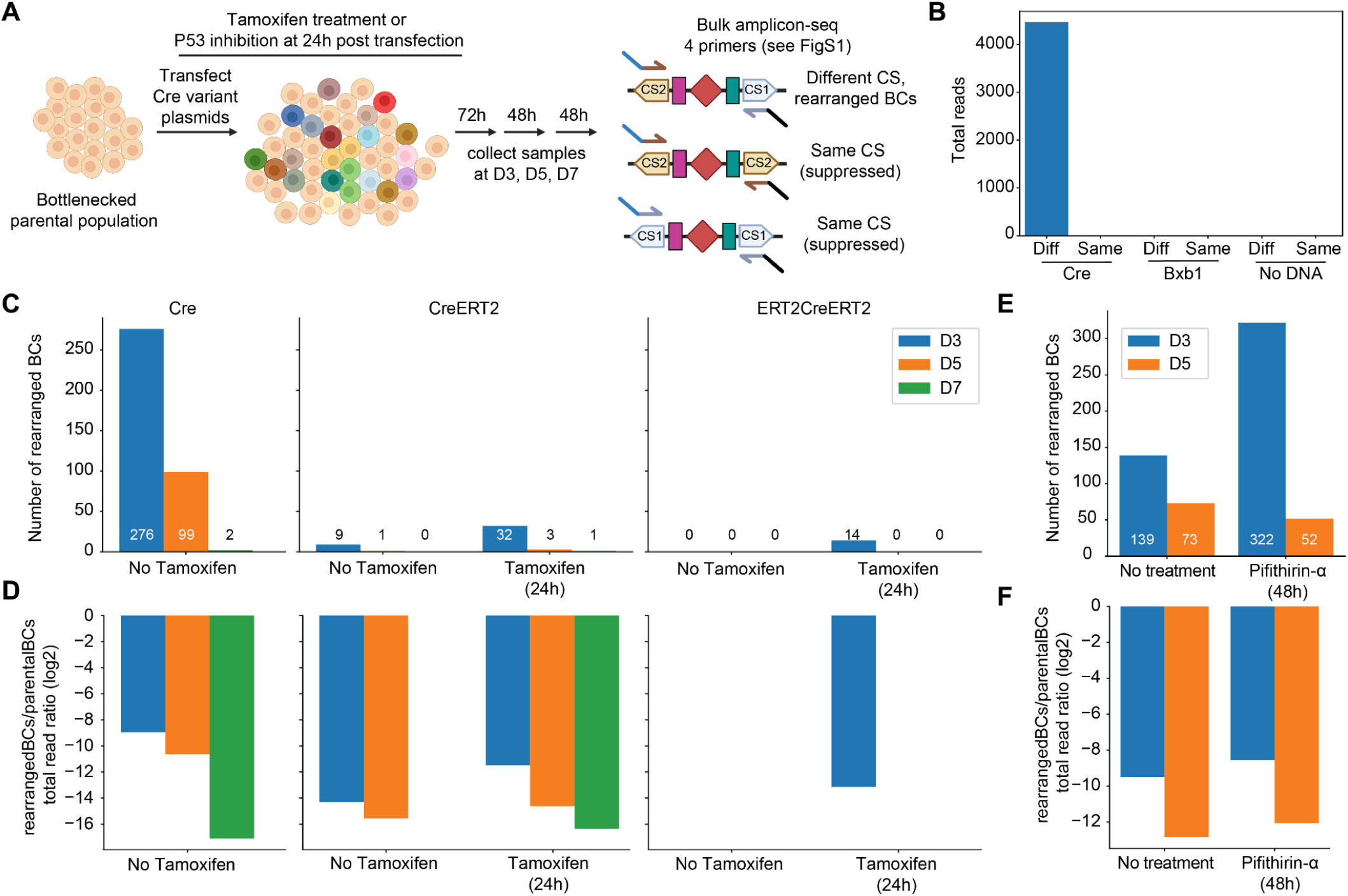
Rearrangements are not stably maintained in the post-Cre induction cell population and cannot be rescued by inducible Cre variants nor by p53 inhibition. **A)** Schematic of the long-term culture experiments with Cre variants or p53 inhibition. The possible products from the 4 primer amplicon-seq strategy (also see **Fig. S1**) are depicted to the right. **B)** Total number of reads from 4 primer amplicon-seq data generated from Cre, Bxb1 or No DNA transfected cells that contain rearranged barcode pairs. Bars are split based on whether the reads contain the same or different (diff) capture sequence on the same molecule. **C)** Number of rearranged barcode (BC) combinations detected at day 3, 5 or 7 post transfection with Cre, CreERT2 or ERT2CreERT2. Cells were either untreated or treated with tamoxifen (0.5μM) for 24 hours. **D)** Log2 ratio of total reads with rearranged BC combinations to parental BC combinations in each sample. **E)** Similar to panel **C**, the number of rearranged BCs detected at day 3 or 5 for Cre-transfected cells with or without p53 inhibitor (Pifithrin-α, 20μM). **F)** Similar to panel **D**, Log2 ratio of total reads with rearranged BC combinations to parental BC combinations for samples with or without p53 inhibitor (Pifithrin-α, 20μM). Data presented in this figure is from one replicate.

**Fig S8.**
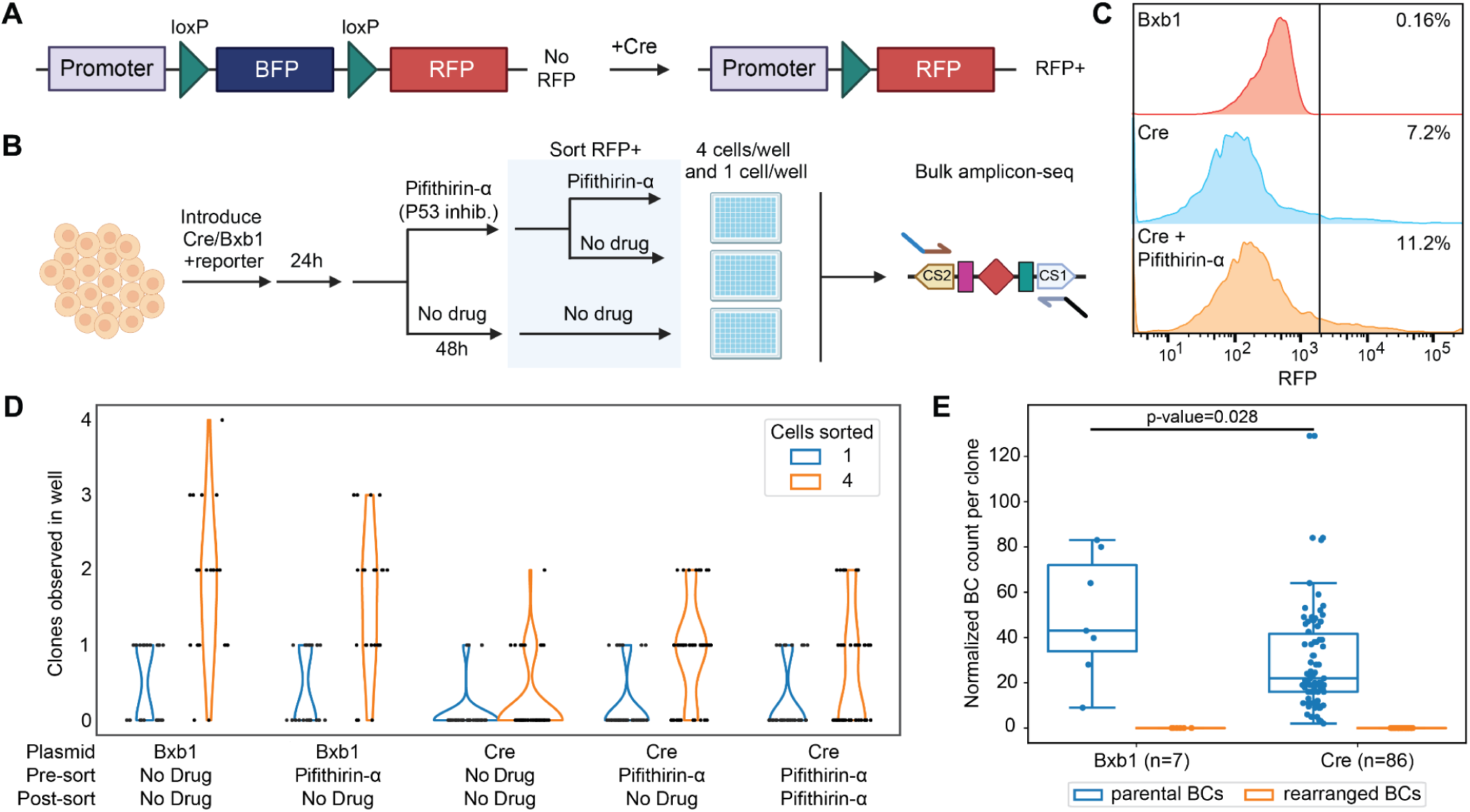
Single-cell sorting does not yield long-lived clones with rearrangements. **A)** The reporter used for this experiment encodes a floxed blue fluorescent protein (BFP) gene which is excised in the presence of Cre to constitutively express a red fluorescent protein (RFP), thus serving as a marker for Cre activity. **B)** Schematic of the single-cell sorting experiment. Cells were transfected with either Cre or Bxb1 recombinase and optionally treated with the P53 inhibitor Pifithrin-α for 48 hours before sorting out either 4 or 1 RFP positive cell(s) into single wells of 96 well plates. Genomic DNA was extracted from clones and barcodes they contained were detected using bulk amplicon-seq. **C)** Flow cytometry traces of cell populations transfected and treated as indicated. The percentage in each panel reflects the RFP positive proportion of the population. **D)** Violin plots depicting the number of clones observed by eye per well 7 days after sorting, separated by the number of cells initially sorted into that well. **E)** Boxplots of the number of parental or rearranged barcode combinations (BCs) observed per well, normalized for the number of clones that were observed in that well. The horizontal solid line indicates the median, the length of the box depicts the interquartile range and the whiskers depict the extent of the distribution minus outliers. Data presented in this figure is from one replicate.

**Fig S9.**
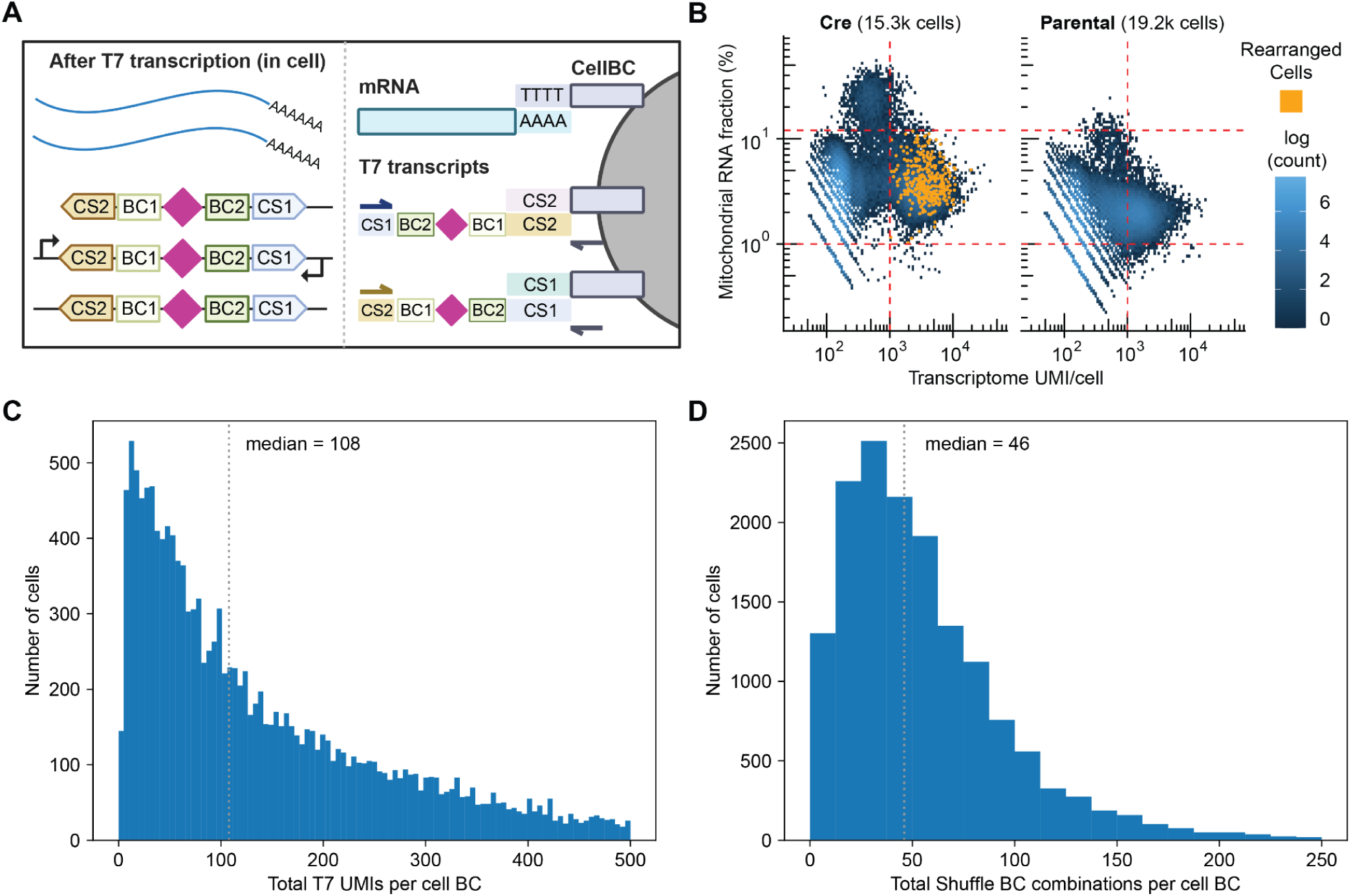
Detection of shuffle cassette barcodes in scRNA-seq data. **A)** After fixation and IVT with T7 polymerase, cells contain both mRNA from endogenous genes as well as RNA from shuffle cassettes. Both of these RNA species can be captured using 10X Genomics gel beads that contain complementary sequences for the polyA on mRNA and capture sequence 1 and 2 (CS1, CS2) found on the T7 derived shuffle transcripts. **B)** Scatter plots of mitochondrial RNA fraction vs. transcriptome unique molecular identifier (UMI) counts per cell detected in the Cre and parental conditions. All cells associated with a rearranged barcode pair at >1 UMI (n=320) are colored yellow. **C)** Histogram of total T7 derived UMIs per cell barcode (BC) with the median value represented by a vertical dotted gray line. **D)** Histogram of total shuffle BC combinations detected per cell in T7 transcripts. Median value is again depicted by a vertical dotted gray line.

**Fig S10.**
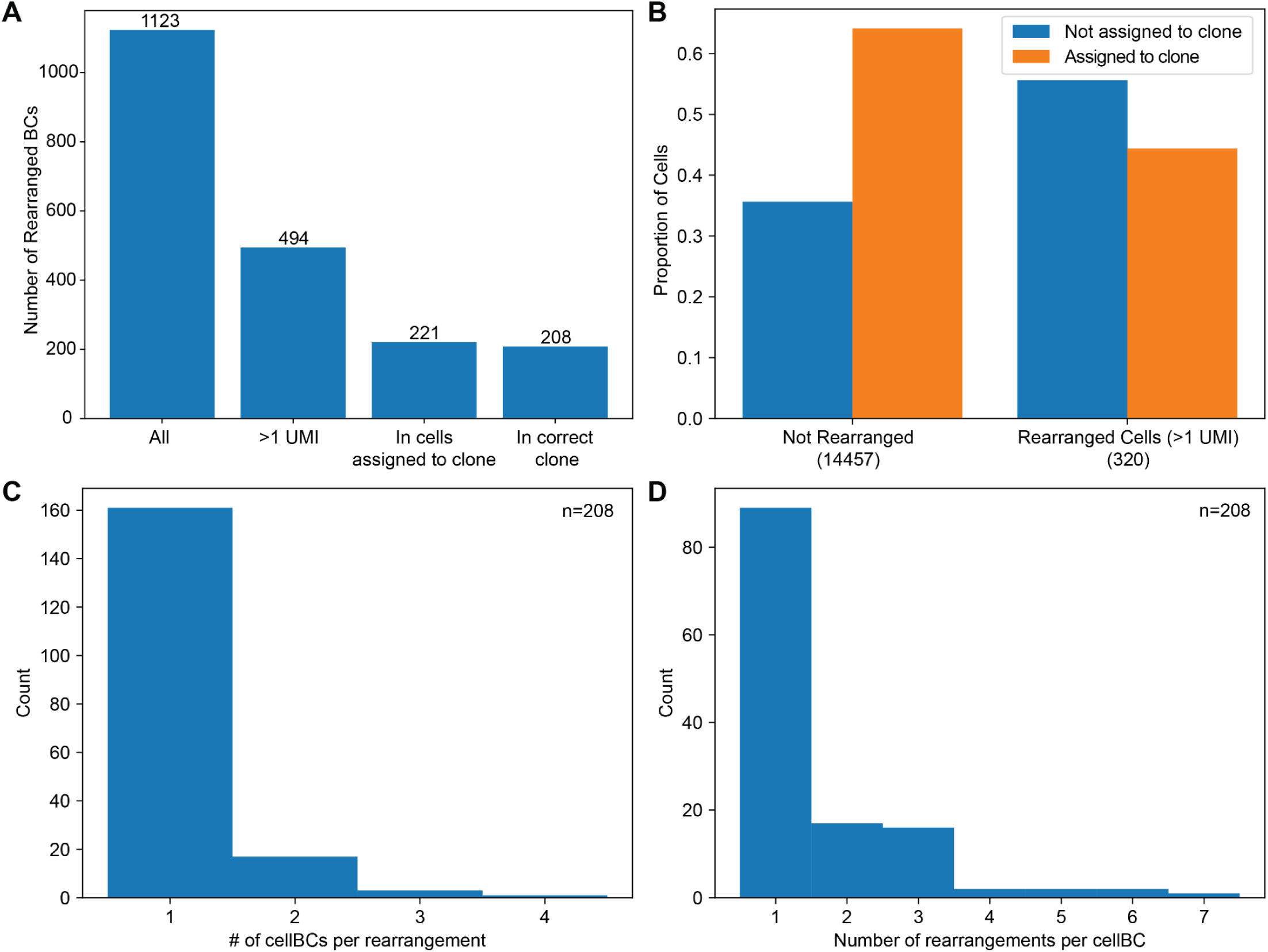
Characteristics of T7-derived barcodes detected in scRNA-seq data. **A)** Number of rearranged BC combinations that are detected in the single cell Cre dataset at successive stages of filtering: all cells, rearrangements detected at >1 UMI, rearrangements at >1 UMI in cells that could be assigned to a clonotype and lastly, rearrangements for which the identity of the rearranged BC pair was congruent with the clonotype assignment. That is, both BCs were detected in the same parental clone. **B)** The proportion of cells that can be assigned to a parental clone for cells that contain a rearrangement or do not. Clonotype assignment is determined by the set of T7 barcodes (BCs) within them, detected with >1 unique molecular identifier (UMI). Cells were considered assigned to a clone if at least 75% of the T7 BCs detected in that cell at >1 UMI belong to that specific clone.**C)** Histogram depicting the number of unique cell barcodes associated with a particular rearrangement. **D)** Histogram depicting the number of rearranged barcodes per cell.

**Fig S11.**
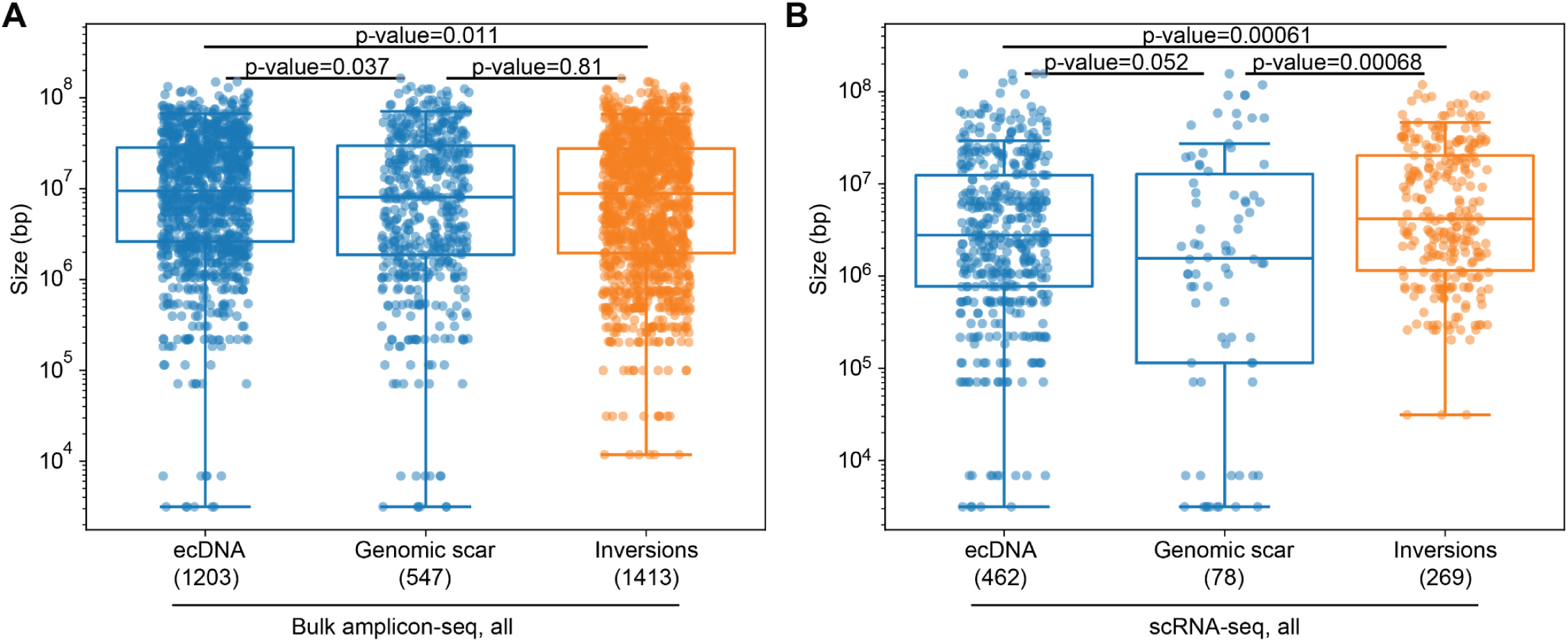
Size distribution of ecDNAs relative to genomic deletions and inversions. **A)** and **B)** Size of each ecDNA, deletion scar and inversion event detected in bulk amplicon-seq data (all rearrangements) and scRNA-seq (all rearrangements) respectively. Depicted p-value is calculated using the non-parametric Mann-Whitney U test. The horizontal solid line indicates the median, the length of the box depicts the interquartile range and the whiskers depict the extent of the distribution minus outliers

**Supplementary Movie 1. Circos plots for the 129 rearranged single-cells detected in the dataset.** Thickness of the line depicting each rearrangement is proportional to its unique molecular identifier (UMI) count in that cell.

## Methods

### Shuffle Cassette Library cloning

The sequence of the shuffle cassette was ordered as a single-stranded oligonucleotide from Integrated DNA Technologies (IDT) with degenerate bases at the appropriate sites to serve as barcodes (**Fig. 1B**). The pool was PCR amplified using Q5 polymerase (NEB M0492S) for 8 cycles with primers that contained overhangs for subsequent cloning. PCR products were run on a polyacrylamide gel (PAGE) and the band at the appropriate size was excised and purified. Purified product was Gibson cloned into a previously described PiggyBac transposon vector (*37*) using standard protocols in a 10 μl reaction (NEB E2621S). 2.5 μl of the Gibson reaction was electroporated into 25 μl of 10-beta electrocompetent *E. coli* (NEB C3020K). The transformed library was grown overnight in 50 mL of liquid Luria Broth (LB) + 50 μg/mL Ampicillin and purified using the ZymoPURE II Plasmid Midiprep kit (Zymo D4200) according to manufacturer’s instructions.

### mESC culture and integration of Shuffle Cassette Library

BL6xCAST mESCs were cultured at 37°C and 5% CO2 on standard tissue culture treated plates coated with 0.1% gelatin in “80/20” medium as previously described (*60*). To integrate the Shuffle Cassette PiggyBac transposon library at high MOI, we reverse transfected 4 μg of library DNA with 200 ng of a PiggyBac Puro-GFP helper plasmid (*54*) and 200 ng of a plasmid expressing the hyperactive transposase hyPBase (*63*) using Lipofectamine 3000 (Thermo L3000001) into 1 million cells. 3 days post transfection, cells were selected with 2 μg/mL Puromycin for 13 days. Surviving cells were bottlenecked to approximately 100 founders by dilution and expanded.

### MOI estimation by qPCR

Genomic DNA was extracted from cells using the Qiagen DNeasy kit. MOI of the shuffle cassette (primers oSP990-oSP993) and Puro-GFP (primers oJBL043-oJBL044) were measured relative to two genomic targets Tfrc (primers oJBL0276-oJBL0277) and Tert (oJBL0280-oJBL0281) which are expected to have 2 copies in these cells (*54*). qPCR was performed using 2X PowerUp Sybr Master Mix (Thermo A25742) with 50 ng of genomic DNA as template per 10 μl reaction with 0.5 μl of 5 μM primer mix. qPCRs were run on a BioRad CFX Opus Real Time PCR instrument in triplicate. Cycle threshold values were averaged across triplicates and copy number was estimated using the ΔCT method relative to each genomic target independently, corrected for the presence of 2 genomic copies and then averaged across the two targets.

### Cre recombinase plasmid transfection and cell population maintenance for shuffle experiments

Recombinase expression plasmids used in this study are pCAG-iCre (Addgene #89573), pCAG-CreERT2 (Addgene #14797) and pCAG-ERT2CreERT2 (Addgene #13777) and pCAG-Bxb1 (pSP0722, modified from Addgene #51271). Between 200,000 and 350,000 cells were reverse transfected with the specific Cre plasmid using Lipofectamine 3000 (Thermo L3000001) in 6-well plates with the exception of the data shown in **Fig. S7E-F** which was generated from a scaled-down version of this protocol performed in 12-well plates in which 300 ng of Cre plasmid was transfected into ∼66000 cells. For experiments depicted in **Fig. S7C-D**, 1 μg of the respective Cre plasmid was transfected. Briefly, DNA and P3000 reagent (2x DNA amount, 8 μl for 4 μg etc.) were mixed with 125 μl of OptiMEM. In a separate tube, Lipofectamine reagent (3x DNA amount, 12 μl for 4 μg etc.) was mixed with 125 μl of OptiMEM. The two tubes were mixed and allowed to sit at room temperature for at least 20 minutes. Transfection mix was added to a gelatinized well of a 6-well plate and cells were added on top in 2mL of medium. Medium was changed every day, and treatments such as 0.5 μM Tamoxifen (Sigma) and 20 μM Pifithrin-α (Sigma) were performed at the specified times. For the Pifithrin-α experiment in which Cre-transfected cells were cultured for longer than 72h, the cell population was split using Accutase (Gibco) and ∼30% of the cell population was transferred to a new plate at day 3. For the inducible Cre variant experiments, cell population was split and 100,000 cells were transferred to a new plate at day 3 and day 5. For all experiments, genomic DNA was prepared from a minimum of 25% of the cell population.

### IVT-seq library construction

An experimental protocol for mapping integration sites using T7 IVT on genomic DNA has been described recently by our group (*37*). In this study, we largely followed this protocol with some minor modifications for each of 2 replicates. The same protocol was followed both for shuffle cassette insertion site mapping from the parental population and for rearrangement call validation post Cre transfection. 300 ng of template genomic DNA purified using the Qiagen DNeasy kit was used as template for all IVT reactions. T7 IVT was performed using the HiScribe T7 High Yield RNA Synthesis Kit (NEB E2040S) in a 60 μl total reaction per sample. Reactions were incubated at 37°C for 16 hours in a thermocycler with the lid set to 50°C. Reactions were treated with Turbo DNase (Thermo AM2238) to remove template DNA according to manufacturer’s instructions and RNA was extracted using Trizol LS reagent (Thermo 10296010). Briefly, each sample volume was normalized to 250 μl with water, 750 μl of Trizol LS reagent was added. Samples were mixed by pipetting and incubated at room temperature for 4 min. 200 μl of chloroform was added and samples were incubated at room temperature for 3 min. Samples were then spun at 12,000 x g for 15 min and the aqueous phase was transferred to a new tube. 1 μl of 5 mg/mL Glycogen was added (Invitrogen) per sample. Next, RNA was precipitated by adding 1 volume of isopropanol. Samples were mixed by inverting and incubated at -80°C for 1 hour and subsequently spun at 21,000 x g for 1 hour at 4°C. RNA pellet was washed with ice cold 80% ethanol and resuspended in 11.5 μl of H2O. Reverse transcription was performed using the SuperScript IV Reverse Transcriptase (Thermo 18090200) with 0.5 uL 100 μM RT primer that contains an 8 base-pair degenerate 3’ end (oSP1012). RNA was initially incubated with 0.5 μL 100 μM RT primer (oSP1012) and 1 μL 10 mM dNTP at 65°C for 5 min and cooled on ice. Enzyme, buffer, RNAse inhibitor and DTT were added and reactions were incubated in a thermocycler at 23°C for 10 min, 50°C for 15 min and 80°C for 10 min, followed by a hold at 10°C.

For replicate 1, 2nd strand synthesis for both top and bottom strands (**Fig. S1B**) was performed in the same reaction with primers oSP1008, oSP1021 and oSP1013. For replicate 2, 2nd strand synthesis for top and bottom strands were performed separately, one with oSP1008 and oSP1013 and another with oSP1021 and oSP1013. Four 50 μl PCR reactions were performed per sample with Q5 Polymerase (NEB M0492S) with the following cycling parameters: 98°C - 3 min; 4 cycles of 98°C - 20s, 65°C - 20s, 72°C - 30s; 72°C - 60s, hold at 4°C. Reactions corresponding to a particular sample and primer pair were pooled. For replicate 1, double-sided size selection (0.5X, 1.1X) was performed with Ampure XP beads (Beckman A63882) on 200 μl of sample and eluted in 50 μl of H2O. For replicate 2, double-sided size selection (0.5X, 1.1X) was performed on 100 μl of sample and eluted in 25 μl of H2O. From each sample, a second PCR was set up with indexing primers using 12 μl of the previous eluate as input. Two 50 μl PCR reactions were performed per sample with Q5 Polymerase (NEB M0492S) with real-time tracking using SYBR green dye with the following cycling parameters 98°C - 3 min; 4 cycles of 98°C - 15s, 65°C - 15s, 72°C - 30s until the curves reached saturation (14-16 cycles). Running these libraries on a D1000 ScreenTape (Agilent) revealed a smear of expected size but also some lower molecular weight products that may dominate the sequencing reaction. To address this, we created 3 equimolar pools: samples from replicate 1, top strand samples from replicate 2 and bottom strand samples from replicate 2. Pools were run on a 6% TBE PAGE gel and DNA between 400 bp and 1000 bp was excised and purified. Library sequencing was performed on an Illumina NextSeq2000 P2 300 cycle kit, with the following read lengths: 106 read1, 10 index1, 10 index2 and 212 on read2.

### Amplicon-seq library construction from bulk samples

As detailed in **Fig. S1**, we employed two different strategies for amplicon-seq library construction: 2-primer (data in **Figs. 3**, **S2C**, **S4**, **S5**, **S6**) and 4-primer (data in **Fig. S7**). The major difference between these two strategies is that either 2 or 4 primers corresponding to the capture sequences were used to generate the PCR product. In the case of the 2-primer strategy, the P5 Illumina sequencing adapter can only come from the primer that binds CS2 and the P7 adapter can only come from the primer that binds CS1. This precludes identification of recombined shuffle cassettes that contain the same capture sequence on both sides of the loxPsym site (**Figs. S1C and S1D**). In the 4 primer strategy, primers with P5 and P7 adapters that bind to both CS2 and CS1 are included in the PCR. Theoretically, this would enable us to detect recombined shuffle cassettes with the same capture-sequence on both sides. However, we do not detect these events even in the case of the 4-primer strategy, probably due to suppressive PCR (**Fig. S1C-D**; **Fig. S7A-B**).

For the 2-primer experiments, we used 250 ng of genomic DNA prepared using the Qiagen DNeasy kit as input. PCR1 (UMI addition) was performed in a 50 μl reaction with Q5 Polymerase (NEB M0492S), with 2.5 μL of each 10 μM primer (oSP1008 and oSP997) with the following cycling parameters: 98°C - 5 min; 4 cycles of 98°C - 20s, 62°C - 20s, 72°C - 30s; 72°C - 60s. PCRs were cleaned up using AmpureXP beads (1X) and eluted in 11 μl of H2O. PCR2 (sample indexing and sequencing adapter addition) was again performed with Q5 Polymerase in a 50 μl reaction using 10 μl of eluate from the previous step as template and 2.5 μL of each 10 μM sample indexing primer. Progression of PCR2 was monitored in real-time using SYBR green dye and reactions were stopped before saturation (usually 15-18 cycles). Cycling conditions for PCR2: 98°C - 5 min; 15 cycles of 98°C - 10s, 65°C - 10s, 72°C - 20s. Reactions were cleaned up with AmpureXP beads (1X) and eluted in 12 μl of H2O. Sample quality was confirmed on a TapeStation D1000 ScreenTape (Agilent), after which reactions were pooled. Sequencing was performed on an Illumina NextSeq2000 100 cycle kit with the following read lengths: 69 read1, 6 index1, 10 index2 and 53 on read2.

For the 4-primer experiments shown in **Fig. S7**, 100 ng of genomic DNA prepared using the Qiagen DNeasy kit as input. PCR1 was performed with a mix of 4 primers (oSP1008, oSP997, oSP1021 and oSP1022) with 0.125 μl of each primer at 100 μM per 50 μl reaction. The rest of the protocol was identical to the 2-primer workflow described above.

### Single-cell sorting and construction of amplicon-seq libraries

Cre reporter (pSP0767 pLV-Flox-BFP-dsRed) was cloned using pLV-flox-dsRed-GFP as a template using Gibson assembly (*64*). 200,000 cells were reverse transfected with 1 μg of Cre or Bxb1 recombinase and 200 ng of the reporter in a 6 well plate per reaction as described above. Two transfections were performed per recombinase. One set of cells per recombinase were treated with 20 μM Pifithrin-α 24 hours post-transfection for a total time of 48 hours. Cells were harvested at 72 hours post-transfection and FACS sorted on the activity of the Cre reporter at single-cell purity into gelatinized 96 well plates containing growth medium with or without Pifithrin-α. FACS data shown in **Fig. S8C** was analyzed using FlowJo.

Clones were allowed to grow out for 9 days before cells were frozen for genomic DNA extraction in 96-well plates. Genomic DNA was extracted in the 96-well format using the Quick-DNA/RNA MagBead kit (Zymo Research R2130) according to the manufacturer’s instructions. Amplicon-seq libraries were constructed from 9.8 μl of template genomic DNA (estimated to be between 10-50 ng) per well using the 4-primer strategy described above. PCR1 was performed in a 20 μl Q5 Polymerase reaction with 0.05 μl of each primer at 100 μM. After 1X clean up with Ampure XP beads and elution in 11 μl of H2O, PCR2 was performed on 10 μl of eluate in a 25 μl Q5 Polymerase reaction with 1.25 μl of each indexing primer at 10 μM. After 18 cycles, 10 μl of each reaction was pooled and purified using a Zymo Research Clean and Concentrate kit. Sample was eluted in 100 μl of H2O, run on an agarose gel and band of the appropriate size was excised and purified (Zymo Research D4007). Libraries were run on an Illumina NextSeq 2000 200 cycle kit.

### Preparation of cells and libraries for single-cell RNA sequencing

300,000 parental cells were transfected in 6-well plates as described above with 1 μg of Cre and 200 ng of the reporter per transfection. 10 individual wells were transfected. At 24 hours post transfection, 5 wells were treated with 20 μM Pifithrin-α for 48 hours total. At 72 hours post transfection, both Pifithrin-α treated and untreated cells were harvested and approximately 500,000 RFP positive cells were FACS sorted into a combined tube based on the activity of the Cre reporter. Untransfected parental cells were harvested and 1 million cells were used as input in parallel with Cre-sorted cells for the protocol. After washing twice with cold 1X PBS (Gibco), cells were resuspended in 400 μl of cold PBS. Cells were fixed with 1600 μl of cold 100% methanol, added dropwise with swirling. Cells were left on ice to fix with gentle swirling to mix every 5 minutes during the incubation. Cells were rehydrated with 4 mL of cold 1X PBS added slowly with gentle swirling of the tube. Cells were spun down and resuspended in 60 μl of PBS. Approximately 60,000 cells and 380,000 cells in total were counted in the Cre and parental samples respectively using a Countess automated cell counter (Thermo) of which all and 100,000 cells in 18 μl of PBS were used for T7 IVT respectively. IVT reactions were set up in 30 μl total volume with the HiScribe T7 High Yield RNA Synthesis Kit (NEB E2040S) with 2 μl of each NTP, buffer and enzyme. Reactions were incubated at 37°C for 1 hour in a thermocycler with the lid at 50°C. Cells were immediately moved to ice and 20 μl of cold PBS was added to each sample. 20 μl and 38 μl (∼40,000 cells) were processed through two independent lanes of a 10x Genomics Single Cell 3’ HT with Feature Barcoding kit.

Transcriptome libraries were prepared as per the manufacturer’s protocol. To prepare libraries from T7 transcripts captured using CS1 and CS2, we started with the supernatant from the clean-up after cDNA amplification of the standard 10x Genomics feature barcoding protocol. Two rounds of PCR were performed with primers specific to the shuffle cassette. In PCR1, oSP997 (CS1-TruSeq2), oSP1022 (CS2-TruSeq2) and oSP1061 (feature-cDNA primer F) primers were used (1.25 μl of 10 μM each) in a 50 μl Q5 polymerase reaction with 5 μl of template. Cycling conditions: 98°C - 45 s; 15 cycles of 98°C - 20s, 60°C - 5s, 72°C - 5s; 72°C - 60s, 4°C - hold. Reactions were cleaned up with AmpureXP beads (1X) and eluted in 15 μl of Qiagen buffer EB. In PCR2, 5 μl of PCR1 eluate was used as input into two 50 μl Q5 Polymerase reactions per sample with sample index primers (1.25 μl of 10 μM each) that add Illumina adapters. Reaction progress was monitored using SYBR green and was stopped before saturation. Cycling conditions: 98°C - 5 min; 9 cycles of 98°C - 10s, 65°C - 10s, 72°C - 20s; 72°C - 60s, 4°C - hold. After a clean up with AmpureXP beads (1X), quality of the libraries was confirmed (prominent single-peak) by running them on a TapeStation D1000 ScreenTape (Agilent). T7 libraries were sequenced on an Illumina NextSeq2000 100 cycle kit with the following read lengths: 28 read1, 6 index1, 8 index2 and 96 on read2. Transcriptome libraries were sequenced on two separate NextSeq2000 100 cycle runs, initially with 28 read1, 10 index1, 10 index2 and 90 on read2 and next with 28 read1, 6 index1, 8 index2 and 96 on read2.

### IVT-seq data analysis for insertion-site mapping in the parental population

Analysis pipeline was based on the pipeline published in ref (*37*). Briefly, reads were demultiplexed using bcl2fastq (v2.20.0.422). Read1 contains the identity of the shuffle barcodes whereas read2 contains the associated genomic sequence. Reads were passed through a custom script (IVTextractBCs.py) to extract barcode sequences based on exact matches to the expected preceding and subsequent bases. The strand of each read (top-CS2 or bottom-CS1) was also assigned at this step. PiggyBac ITR sequences were trimmed from read2 using cutadapt (v2.5) with the following parameters: -cores=4 --discard-untrimmed -e 0.2 -m 10 -a CCCTAGAAAGATA (*65*). Trimmed reads were mapped to mm10 reference genome using bwa mem (v0.7.17) with -Y option (*66*). SAM files were sorted using samtools (v1.9) and filtered out reads that do not align to known PiggyBac insertions sites (TTAA) or align to several locations (contain XA:Z flag) using a custom script (align_filter.py) (*67*). Filtered SAM files were converted to BED format using the sam2bed tool in bedops (v2.4.35) (*68*).

Bedtools (v2.29.2) intersect was used with the -loj -wa -wb -filenames -sorted options to extract those alignments overlapping with a known variant between the BL6 and CAST alleles from the Sanger Mouse Genome database (*39*, *69*). A custom python script (cleanup_sort_variantcall_update.py) was used to parse the CIGAR string of each alignment intersected BED file to assign each read to each of the following categories: BL6 (read overlaps with variant and sequence of alignment at that position matches the reference allele), CAST (read overlaps with variant and sequence of alignment at that position matches variant allele) and ‘no-Variant’ (read does not overlap with variant). Inconclusive alignments that contained some incongruence in allele assignment were discarded. The position of each alignment was determined by the genomic strand it mapped to. If strand is +, position is the end of the alignment and conversely, if strand is -, position is the start of alignment.

Next, another custom python script (ivt_clustered_groupcollapse_iterable.py) was used to collapse alignments into groups based on: 1) a shared chromosome and position and 2) a shared set of barcodes extracted from read1. Alignments at a given position were clustered together if their associated barcodes were within a Levenshtein distance of 6. Within each cluster, the most common values for barcode1, barcode2, strand of alignment to genome (+ or -) and shuffle cassette strand (top-CS2 or bottom-CS1) were assigned as the representative values for that cluster. Each cluster was assigned to an allele based on the most common allele value within, without taking into account the number of no-Variant assignments within that cluster. Total read count (number of alignments per cluster), total read1 UMIs (coming from forward primer of second strand synthesis), total read2 UMIs (first 8 bp of genome coming from the degenerate 3’ end of the RT primer) and total number of unique alignment lengths per cluster were also determined.

To define the list of parental insertions, the outputs from the previous step from replicate 1 and replicate 2 were merged based on shared position, barcode and strand values. Within this merged dataset (n=31,817), for each cluster of alignments, allele value was assigned based on the allele value in replicate 1 and replicate 2, without considering no-Variant assignments. In the case that replicate 1 and replicate 2 allele assignments were incongruent, allele for that cluster was assigned as ‘inconclusive’. Clusters with <6 reads and <3 unique lengths aligned in replicate 2 of the parental sample were filtered out leaving 22,158 clusters. Clusters containing barcode combinations that did not map uniquely to a genomic location were then filtered out to leave 12,695 clusters. Within this set, bonafide insertions were defined by: 1) a pair of read clusters whose alignment position differs by exactly 4 bp, 2) the first cluster in the pair maps to the - strand and the second cluster maps to the + strand, 3) the pair does not encode the same shuffle cassette strand (top-CS2 or bottom-CS1); and 4) the barcodes detected in each member of the pair are reverse complement of one another. This resulted in 5,145 pairs of clusters identified as shuffle cassette insertions.

Within this set, the allele value for each insertion was assigned as inconclusive in the case that the allele assignment for the pair of read clusters that make up the insertion were incongruent. In the case that one of the clusters were marked as no-Variant, the allele of the other cluster in the pair was considered the allele of that insertion. Insertions were merged with amplicon-seq data from the parental cells based on shared barcode pairs. Only those insertions whose barcodes could be detected in the amplicon-seq data from the parental cells (see more details below) with >50 reads and >30 UMIs were kept (n=5,095). There were a small number of remaining sites (n=7) that had more than one pair of barcodes called at that site. By manual observation, these seemed to arise from a clustering artifact and ought to have been collapsed into one cluster. We arbitrarily chose the row that had the higher value in the number of unique lengths aligned in rep2 on the left side of the insertion at these positions for the final set (n=5,088).

To generate the visualization in **Fig. 2A**, alignments were visualized in IGV 2.16.1 (*70*) and BED files containing insertions assigned to each allele were loaded separately as tracks. The visualizations in **Fig. 2D** and **Fig. S3** were generated using the ChIPseeker package in R (*71*). Other plots were made using a combination of matplotlib (3.8.1) and seaborn (0.13.0) libraries in Python.

### Amplicon-seq analysis and rearrangement calling

Reads were demultiplexed using bcl2fastq (v2.20.0.422). Reads were passed through a custom script (read_extract_iterable.py or read_extract_iterable_4primers.py) to extract barcode sequences and UMIs based on exact matches to the expected preceding and subsequent bases. In the case of 4-primer amplicon-seq, the strand of each read (top-CS2 or bottom-CS1) was also assigned at this step. Total number of reads per barcode pair was taken as the readcount and the total number of unique UMIs detected was taken as the UMI count (df_group.py or df_group_4primers.py). In 4-primer amplicon-seq, the same shuffle cassette can result in two distinct amplicons. We collapsed reads coming from the same shuffle cassette based on the shared set of barcodes and summed up the read counts and UMI counts. Those barcode pairs where we did not detect both types of amplicons were discarded. In all cases, the barcode closest to CS2 was named barcode1 and the barcode closest to CS1 in the shuffle cassette was named barcode2. Read and UMI counts were normalized for sequencing depth. For the parental barcode set, normalized read and UMI counts were averaged across 4 replicates and used for defining the bonafide set of insertions as detailed in the section above.

For the plot in **Fig. S7B**, we looked for amplicons with shared barcode pairs that contained the same capture sequence. To eliminate confounding through errors in PCR or sequencing, we restricted our search to those amplicons containing barcodes which were both found in the bonafide list of parental insertions. We also eliminated those amplicons that contained the same barcode (that is, within Levenshtein distance of 6) on both sides of the loxPsym site. CS1-CS1 and CS2-CS2 amplicons were not readily detected in our data, presumably due to suppressive PCR (**Fig. S1**; **Fig. S7B**).

For the amplicon-seq data generated from single-cell sorted clones (**Fig. S8**), wells with fewer than 100k reads were discarded. Barcode sequences were extracted and read/UMI counts per barcode pair were determined as above. The set of barcodes associated with each well were determined as the barcode pairs whose readcounts were >1 standard deviation above the mean readcount for barcode pairs detected in that well using the zscore function in the scipy.stats library. Analysis was then restricted to those barcode pairs that were present in the bonafide parental insertion list (n=5088). Barcode pair count per well was normalized for the number of clones observed in that well by eye.

Rearranged barcodes were identified by comparing the set of identified barcode pairs in the dataset to the bonafide list of parental insertions. Both barcodes in the rearranged pair were required to be in the bonafide list but were not found together in the parental cells. To remove any artifacts caused by PCR chimeras or errors, each rearranged barcode pair was required to be present at >1 UMI. The nature of each rearrangement denoted by a rearranged barcode pair was inferred based on the position and orientation of the parental insertion sites (**Fig. 1**; **Fig. S1**). We first determined that inter-homolog translocations were rare (recombination between shuffle cassettes on the same chromosome but assigned to different alleles). Therefore, we parsimoniously identified deletions as those rearranged barcode pairs between shuffle cassettes on the same chromosome that were inserted in the same orientation. In a similar manner, inversions were identified as those rearranged barcode pairs between shuffle cassettes on the same chromosome that were inserted in the opposite orientation. The size of the rearrangement was calculated as the difference between the position of the two original insertion sites. Translocations were those rearranged barcode pairs found between shuffle cassettes on different chromosomes.

Deletions could be further classified as coming from the genomic copy or the extrachromosomal circle based on the barcodes that were detected (**Fig. 1**; **Fig. 2D**). For example, let us consider the case of a deletion between two shuffle-insertions X and Y at pos N and pos N+100 on a chromosome, with ‘top-CS2’ orientation (CS2 found closest to the left of the chromosome). The ecDNA would contain the CS2 barcode of insertion Y and and CS1 barcode of insertion X and the genomic copy would contain the CS2 barcode of insertion X and and CS1 barcode of insertion Y. In the case of deletions between two ‘bottom-CS1’ insertions with (CS1 found closest to the left of the chromosome), the opposite would be true.

All Circos plots were made using the pyCircos library (*72*). The remaining plots were made using a combination of matplotlib (3.8.1) and seaborn (0.13.0) libraries in Python.

### Rearrangement call validation using IVT-seq data

IVT-seq libraries were constructed using the same genomic DNA samples used to prepare amplicon-seq libraries from Cre-transfected samples. The data was pushed through the same analysis pipeline as described above for IVT data from parental cells, until the collapsing of alignments into groups based on their shared barcodes and positions. For each rearrangement detected in the amplicon-seq data, we asked whether there was at least one transcript detected in the IVT-seq data from that sample that supported the rearrangement call (**Fig. 3D**). The fraction of rearrangements supported in each IVT replicate from each sample were plotted using matplotlib (3.8.1) library in Python.

### Single-cell data analysis

#### Initial pre-processing and quality filtering

10x Genomics 3’ gene expression (transcriptome) libraries were processed using cellranger-6.0.1 count function (with reference refdata-cellranger-mm10-3.0.0). Resulting raw count matrices were converted to a Seurat (v4.3.1) (*73*) object using functions Read10X and CreateSeuratObject (options: min.cells=3, min.features=50). The mitochondrial fraction was computed and cell barcodes with >1000 transcriptome UMI/cell and with 1-12% mitochondrial fraction (**Fig. S9B**) were retained. Scrublet 0.2.3 (*74*) was run on filtered cells, and cells with doublet score <0.4, leaving ∼15.3k (Cre treated) and ∼19.2k (Parental) high-quality cells for downstream analysis.

We note that a partial wetting failure in the parental sample is the probable cause of less clean separation of bonafide encapsulated cells from ambient RNA containing droplets (**Fig. S9B**). The same quality thresholds were nevertheless used to identify cell barcodes from the parental population for downstream analyses. The Cre-treated sample displayed a cleanly separated cell population in the total transcriptome vs. mitochondrial fraction plane, indicating good cellularity despite fixation and IVT treatment to generate the T7-BC prior to emulsion and library generation.

#### Generating T7-shuffle BC count matrix

To obtain the BC-by-cell-count matrix, T7-BC sequencing data was first processed using cellranger-6.0.1 count to perform error correction on the cell barcodes. The unmapped reads with successfully error-corrected UMIs were selected from the sorted BAM file output, and barcodes sequences were extracted for each read from the shuffle cassette by looking for matches for constant surrounding sequences (CS2 captured IVT RNA: BC1 from TGAGC(.{20})ATAAC in positions 15 to 49 of the read & BC2 from GTTAT(.{20}) in positions 69 until the end; CS1 captured IVT RNA: BC2 from AAAGC(.{20})ATAAC in positions 15 to 49 in the read and BC1 from GTTAT(.{20}) in positions 69 until the end), and joined to the error-corrected cell barcode and umi from the read. Reads counts and total set of UMIs for all cellBC/BC1/BC2/capture-orientation combinations were then tallied, discarding likely chimeric UMIs (taken to be UMIs for which the proportion of reads associated to a given BC1/BC2/capture-orientation all other BC1/BC2/capture-orientation in the specified cell barcode falls below 0.2). The number of error-corrected UMIs for a given BC1/BC2/capture-orientation was then taken as the number of connected components in a graph created by connecting all UMIs associated with that BC1/BC2/capture-orientation combination with a Hamming distance≤1.

#### De novo identification of clonotypes

Similarly to previous works (*54*, *75*, *76*), we leveraged high MOI (many T7-BC pairs per cell) and clonal nature of the population to identify clonotypes (defined as the full set of BC pairs within a clone) directly from the single-cell data T7-BC data, working under the assumption that co-detection of T7-BC pairs should only happen from barcodes within the same clone.

We first summed UMI counts in the same cell with the same BC1/BC2 pair but captured from different capture sequences. We then subsetted the T7-BC UMI counts to those originating from the set of quality filtered cells (Cre-treated) as described above, and retained barcode pairs with >= 3 UMI per barcode per cell. To further remove cells and barcode pairs with little signal for the purpose of *de novo* clonotype identification, were removed any cells with <20 total T7-BC UMIs (after the >=3 UMI/BC per cell thresholding) and T7-BC pairs with <20 total UMI across all cells, leaving ∼8.6k high representation T7-BC pairs and ∼11k cells. We then constructed a count matrix and used Seurat v4.3.1 to perform dimensional reduction (NormalizeData with normalization.method=”RC”, scale.factor=10000, FindVAriableFeatures with selection method=”vst”, nfeatures=length(T7_BC), RunPCA on the top 100 PCs with identified variable features, FindNeighbors with k.param=10, and FindClusters at resolution=1). The resulting 146 communities were used to identify the putative set of T7-BC per cluster.

To do so, T7-BC representation across all cells assigned from clustering in the BC space was calculated as the proportion of cells in the cluster with the barcode pair detected. These proportions were then rank ordered, and a heuristic threshold was used to demarcate barcode pairs associated with a clone: the threshold was set as the fold-change in representation from rank n BC pair to rank n+1 barcode pair became higher than 1.5, empirically selected as a reliable marker of the inflection point in the distribution. The resulting set of putative clonotypes were further filtered by retaining only clonotypes for which the maximum detection fraction (from the top T7-BC pair for that clonotype) was above 0.5, and in which >6 T7-BC pairs were detected. In addition, subsetted T7-BC count matrix for each putative clonotype (with only assigned cells and contained T7-BCs) was inspected, and any clonotype corresponding to a clear doublet (split in the matrix in two blocks) was not retained for downstream analysis (3 putative doublet clonotypes on round 1). As a final quality control step, because this procedure tends to redundantly create multiple clusters for the same clonotype (depending on the resolution parameter), we computed the Jaccard index (on the set of T7-BCs) for each identified clonotype pair. The distribution of Jaccard indices was very strongly bimodal, with the majority of pairs displaying 0 overlap, with a small minority (12 pairs) with Jaccard index of >0.95, suggesting identical underlying clonotypes. A graph was created with nodes corresponding to putative clonotypes and connected if their Jaccard index was >0.1. The intersection of the T7-BC from the connected clusters (mostly singletons) was then taken as the clonotypes. All in all, this led to 108 high confidence *de novo* clonotypes from round 1 (see below for round 2), with a mean MOI of 60.2.

We note that while re-arranged barcodes could be considered as adding a possible source of noise in this analysis, we have found none of the barcode pairs to correspond to chimeric pairs not found in the parental population as determined from bulk sequencing of the parental barcode pairs.

#### Comparison of de novo and bulk-derived clonotypes

As a control for the validity of our *de novo* clonotypes, we directly compared to clonally expanded cells sampled from our bottlenecked population, from which barcode pairs were extracted in bulk by PCR amplification and sequencing from genomic DNA extracted from separately expanding pools, which resulted in a set of 49 ‘bulk’ clonotypes after Jaccard index merging of identical barcode sets. For every pair of clonotypes (both ‘bulk’ and *de novo* from the single-cell data), the Jaccard index was calculated as before. This again resulted in a nearly perfectly bimodal distribution, with the overwhelming majority of pairs completely disjoint, with a small set of 38 pairs with Jaccard index>0.05 with near perfect overlap (mean Jaccard index = 0.961). This confirmed that we were able to capture a large proportion (38/49) of the clonotypes from *de novo* reconstruction from the single-cell data, and that the identified *de novo* clonotypes were highly congruent to the ground truth set constructed from bulk monoclonal derived barcode sets. Given uneven sequencing saturation of the bulk-derived clonotypes, our first set of clonotypes for assignments of cells to clone was taken as the union of *de novo* round 1 clonotypes with (non-overlapping) bulk clonotypes, or 119 clonotypes.

#### Iterative round of assignment and mapping to identify additional clonotypes

In order to comprehensively identify clonotypes, we performed an iterative approach whereby cells were first assigned to clonotypes, as described using the 119 round 1 set described above. Any assignable cell was then removed from the *de novo* pipeline described previously, and the process repeated. Doing so generated an additional set of 24 high confidence clonotypes, for a combined set of 143 high-confidence clonotypes for final assignments.

#### Single-cell assignment of cells to clonotypes

To obtain a more sensitive assignment of cells to clonotypes (in contrast to the clustering in a dimensionally reduced barcode space used for *de novo* clonotype calling as described above), we compared all the T7-BC detected (considering only events with >= 2 UMI per barcodes pairs per cell, after summing pairs captured from both CS1 and CS2) in all cells to to the set of high-confidence clonotypes. In order not to bias against re-arranged cells for the purpose of assignment to clones, barcodes were considered separately for this analysis (and found to be highly congruent, such that the assignment was performed from barcode 1). Specifically, for each cell-clonotype pair, the fraction of cell-detected barcodes belonging to the clonotype (precision) was recorded, in addition to the fraction barcodes from the clonotype recovered in the cell (recall). Across all cells, the clonotype with the highest precision (similar result if selecting on recall) was retained. The resulting distribution of top scoring clonotype precision/recall assignments displayed a peak around 0.6 top_recall and 0.9 top_precision. Cells in that plane were considered assignable to a clone with high confidence if top_recall>0.1 and top_precision>0.75. To further remove the possibility of doublets, any resulting assigned cells with second-top assigned clones showing recall > 0.1 were removed from the set of high-confidence assignments. In the end, 63.6% (∼9.4k/∼14.8k) of cells with at least one BC pair with >= 2 UMI) of cells were assignable to a clonotype. Of the remaining cells, 2.6k displayed low capture (recall < 0.1) either as a result of missed clonotypes from our reconstruction procedure or from low levels of IVT for the T7-BC generation. The remaining cells, where the set of detected T7-BCs are not predominantly from a single clonotype, either originated from droplets encapsulating doublets or large quantities of ambient RNA.

#### Precision-recall analysis for T7-shuffle BC detection

In order to assess which thresholds to use to identify high-confidence detections of rearrangement events, we considered high-confidence assigned cells to clonotypes, and determined precision and recall values for varying UMI thresholds. At each UMI threshold, the precision and recall values were averaged over all high-confidence cell-to-clonotype assignments (**Fig. 4D**) The curves displayed a sharp increase at 2 UMI (1 UMI mean precision 0.53 to 2 UMI mean precision of 0.94) with marginal decrease in recall (0.58 to 0.48). While this corresponds to an upper bound on performance given that only cells with good capture would be assignable in the first place (we confirmed similar performance with a looser threshold assignment of 0.75 precision on the top scoring clonotype), we consider the assessment as providing solid empirical ground for taking the threshold of 2 UMI as associated with a <5% FDR.

#### Rearrangement calling and visualization from single-cell data

From the set of T7-BCs identified in the single-cell data, rearrangements were called as described above for bulk amplicon-seq. Out of the 1123 rearranged BC pairs identified in the dataset, 494 were present at >1 UMI, of which 221 were found in cells assigned to a clonotype as described above. For this set of rearrangements we asked if the pair of BCs participating in the rearrangement originated from the same clone that the cell was assigned to. 208/221 rearrangements were consistent with the clone of the cell they were detected in. The remaining pairs of rearranged BCs come from the same clone but not the clone that the cell was assigned to, indicating that these likely arose from ambient RNA contamination. The subsequent analysis focused on the validated set of 208 rearrangements. Circos plots were made using the pyCircos library and arranged into a movie with iMovie. Other plots were made using a combination of the matplotlib (3.8.1) and seaborn (0.13.0) libraries in python.

## List of Supplementary Tables

**Supplementary Table 1**. List of primers and their sequences used in this study.

**Supplementary Table 2**. List of parental shuffle cassette insertion positions.

**Supplementary Table 3**. All rearranged barcodes detected in 2-primer bulk amplicon-seq at 72h post Cre treatment (data used in Figs. 3, 5, S4-6, S11).

**Supplementary Table 4**. All rearranged barcodes detected in both technical replicates of 2-primer bulk amplicon-seq at 72h post Cre treatment (data used in Figs. 3, 5, S4-6, S11).

**Supplementary Table 5**. All rearranged barcodes detected in the Cre sample of the 10X single-cell RNA seq experiment (data used in Figs. 4, 5, S10-11).

**Supplementary Table 6**. Rearranged barcodes detected in cells assigned to clone at >1 UMI in the 10X single-cell RNA seq experiment (data used in Figs. 4, 5, S10-11).

**Supplementary Table 7**. List of clonotypes and associated barcodes identified from single-cell RNA sequencing and amplicon-seq from single-cell sorted clones.

**Supplementary Table 8**. Assignment of cells to clonotypes based on the set of T7-derived barcodes detected within them.

**Supplementary Table 9**. Details of all sequencing libraries presented in this study, as well as description of processed file and description of wells from the single-cell sorting experiment described in Fig. S8.

## Acknowledgements

We thank Ran Brosh for sharing plasmids and the NYU CEGS group, members of the Shendure lab, Danny Miller and Chia-Lin Wei for helpful discussions. We also thank Choli Lee for assistance with FACS sorting. We acknowledge the use of Biorender.com to generate schematics for many figures in this manuscript.

## Funding

This work was supported by research grants including NIH DP5OD036167 to S.P., NHGRI CEGS subward RM1HG009491 to S.P., NHGRI R01HG010632 to J.S, and funds provided by the Brotman-Baty Institute for Precision Medicine. J.B.L. is a fellow of the Damon Runyon Cancer Research Foundation (DRG-2435-21). J.S. is an investigator of the Howard Hughes Medical Institute.

## Author contributions

S.P. conceptualized the *Genome-shuffle-seq* approach with help from J.S., J.B.L, R.D., J.K. and X.L. S.P. and R.D. performed all experiments. S.P. and J.B.L analyzed data. J.B.L, D.L and X.L piloted the capture of T7 IST transcripts within single-cells using the 10x Genomics platform. S.P, J.S and J.B.L wrote the manuscript with input from all authors.

## Competing interests

J.S. is a scientific advisory board member, consultant and/or co-founder of Prime Medicine, Cajal Neuroscience, Guardant Health, Maze Therapeutics, Camp4 Therapeutics, Phase Genomics, Adaptive Biotechnologies, Scale Biosciences, Sixth Street Capital and Pacific Biosciences. The other authors declare no competing interests.

## Data/Code Availability

Analysis pipelines and plasmid sequences can be found at: https://github.com/sudpinglay/genome-shuffle-seq

Raw sequencing data and processed files can be found at: https://krishna.gs.washington.edu/content/members/pinglay/sud/genome_shuffle_geo/ Data will be deposited in the Gene Expression Omnibus (GEO).

## Notes

### Summary of Updates

Some minor revisions to text, figures and methods. Included links to raw data and analysis scripts.

